# Met32 governs transcriptional control of sulfur metabolic flexibility and resistance to reactive sulfur species in the human fungal pathogen *Candida albicans*

**DOI:** 10.1101/2025.05.12.653471

**Authors:** Anagha C.T. Menon, Faiza Tebbji, Azadeh Alikashani, Antony T. Vincent, Adnane Sellam

**Author notes:** Address correspondence to Adnane Sellam.

## Abstract

Although considerable advances have been made in understanding the metabolic machinery that enables bacteria to utilize sulfur sources, many aspects of this process remain understudied in fungi. To explore the genetic circuit by which the highly prevalent human opportunistic yeast *Candida albicans* controls sulfur utilization, we characterized the transcriptional landscape associated with sulfur starvation in this fungus. We identified many desulfonation enzymes that were differentially modulated and showed that Jlp12, a sulfonate/α-ketoglutarate dioxygenase, was critical for the utilization of different sulfur sources found in many niches of the human host. We also uncovered that the Zinc-finger transcription factor Met32 acts as a master regulator, that modulates genes of sulfur utilization including Jlp12. Importantly, we found that *C. albicans* Met32 exclusively regulates sulfur utilization genes, while in the *Saccharomyces cerevisiae* lineage, it controls methionine biosynthesis. This work also identified Seo13 as the first major facilitator superfamily transporter in fungi that transports the alternative sulfur source glutathione, under the direct control of Met32. Furthermore, we showed that Met32 modulates *C. albicans* tolerance to sulfite excess by tuning the basal transcriptional level of the superoxide dismutase Sod1. This underscores the dual role of Met32 in the breakdown of sulfur-containing metabolites and the neutralization of the resulting reactive sulfur species (RSS). Our study delineates a new mechanism by which fungal pathogens utilize sulfur sources and neutralize RSS and underscores its importance in fungal fitness *in vivo*.

**Importance:** *Candida albicans* is the most prevalent fungal colonizer of humans and it is also the first cause of disseminated fungal infections leading to a high mortality rate. The ability of this yeast to metabolize a plethora of carbon and nitrogen sources inside the host is a critical asset for both the commensal and the pathogenic lifestyles of this yeast. Thus, these pathways represent attractive targets for antifungal therapy. While sulfur is an essential nutritional element for all living organisms, its contribution to fungal virulence remains understudied. Here, we describe new players of sulfur utilization metabolism in *C. albicans* and underline their importance in supporting fungal virulence. This work emphasizes the significance of targeting sulfur metabolic flexibility to manage fungal infections.

## Introduction

The ability of fungal pathogens to utilize a vast array of nutrients in the different colonized niches of the human host, also termed metabolic flexibility, is critical for their growth and persistence. While the vast majority of works investigating the contribution of metabolic flexibility to fungal virulence have focused mainly on carbon and nitrogen (1, 2), sulfur has received only little attention. Sulfur is an important trace element that is essential for the synthesis of the proteogenic amino acids methionine and cysteine, in addition to many sulfur-containing metabolites such as glutathione (GSH), thionucleosides and enzyme co-factors (3, 4). Given the importance of sulfur in biological systems, the ability of fungal pathogens to uptake and utilize both organic and inorganic sulfur-containing molecules might be of high significance for their fitness. The importance of sulfur metabolism in fungal virulence is supported by works in the airborne opportunistic fungus *Aspergillus fumigatus*, where different sulfur metabolic routes including methionine biosynthesis and the production of the sulfur-containing mycotoxin, gliotoxin, were shown to be essential for host infection (5–8). In *Cryptococcus neoformans,* the causative agent of fungal meningitis, genetic inactivation of the bZip transcription factor Cys3 that controls sulfur assimilation and methionine biosynthesis led to attenuated virulence (5, 9). In the prevalent human pathogen *Candida albicans*, the causal role for sulfur metabolism in fungal pathogenicity remains understudied (10). Recent work emphasized the importance of methionine uptake by the high affinity permease Mup1 in the ability of *C. albicans* to infect the host (11). Overall, these studies underscore the importance of sulfur metabolism in fungal virulence and its potential as target for antifungal therapy. Druggability of fungal sulfur metabolism is also supported by the fact that many proteins of this pathway are not conserved in the human host (12).

The core sulfur metabolic machinery has been extensively deciphered in the budding yeast *Saccharomyces cerevisiae* (13, 14). Yeast cells uptake sulfate (SO4^2-^), the preferred sulfur source for fungi, from the extracellular environment using high affinity transporters Sul1 and Sul2 (13, 14). Intracellular sulfate is then assimilated through different sequential reactions to sulfide (H_2_S) that is incorporated into a carbon chain to generate precursors that feed the methionine and cysteine biosynthesis (13). Yeast cells are also able to uptake sulfur-containing metabolites such as methionine and cysteine from their environment using both specific transporters (Mup1 and Mup3) and general amino acid permease (Gap1) (15). A major transporter of both inorganic sulfur sources and sulfonates was recently identified in *S. cerevisiae* underscoring the large flexibility of fungal cells toward sulfur source utilization (16). Sulfonates such as taurine are alternative sulfur sources that are degraded by the _α_-ketoglutarate dioxygenase Jlp1 to generate sulfite (SO ^2^) that is assimilated by *S. cerevisiae* cells (17). At the transcriptional level, genes of sulfur assimilation and methionine/cysteine biosynthesis are tightly monitored depending on the availability of sulfur sources (18). Met4 is a bZIP transcriptional activator that is a central modulator of sulfur metabolism in *S. cerevisiae* and controls the majority of genes of the sulfur metabolic network (13, 19). Met4 lacks DNA-binding domain and depends, for its recruitment to promoters, on the interaction with three transcription factors: Cbf1, Met31/32 and Met28 (20). Conversely, these transcription factors rely entirely on Met4 for transcription activation and serve as a platform for Met4 recruitment (18). The two functionally redundant Zinc-finger transcription factor paralogs, Met31 and Met32 bind to CTGTGGC motif, while the bHLH transcription factor Cbf1 recognizes CACGTG (19, 21, 22). When sulfur-containing metabolites such as S-adenosyl methionine (SAM) and cysteine are abundant in *S. cerevisiae* cells, Met4 is inactivated by ubiquitination that is mediated by the ubiquitin ligase Met30 leading to the inactivation of the Met4-Met31/32-Cbf1 transcriptional module (18).

Overall, the sulfur assimilation and methionine biosynthetic pathways are well conserved in *C. albicans* with few unique features (23). For instance, genetic inactivation of *MET15* encoding the O-acetylhomoserine-O-acetylserine sulfhydrylase, one of the enzymes of the sulfur assimilation route, does not lead to sulfur auxotrophy as seen in *S. cerevisiae* (24). Furthermore, the cobalamin-independent methionine synthase Met6 was found to be essential while it is not in *S. cerevisiae* (25). Recent studies have shown that, while the role of Met4 as a central regulator of methionine biosynthetic genes (*MET*) is conserved, Met32 function has diverged as the *met32* mutant has no auxotrophy for methionine and the Met32 motif was not enriched in the promoter of *MET* genes (26). The role of *C. albicans* Cbf1 in sulfur uptake and activation of *MET* genes is conserved as Cbf1 binds to the promoters of sulfur and methionine transporters in addition to *MET* genes (26–28).

In the human host, sulfur metabolites are present under different organic (e.g. proteins/peptides, methionine, and cysteine) or inorganic forms (SO_4_ ^2-^, SO_3_ ^2-^ and H_2_S). In the GI tract, which is the large reservoir of *C. albicans*, organic sulfur-containing metabolites include free cysteine and methionine, sulfomucin and taurine-conjugated bile acids while the inorganic forms are represented mainly by dietary sulfates or sulfide produced by gut microbiota (29, 30). So far, studies on sulfur utilization in pathogenic fungi were mainly focused on pathways using inorganic sulfur ions or sulfur amino acids and have underestimated how other alternative sulfur sources such as taurine, the highly abundant intestinal organosulfonate, are assimilated (31). Intestinal bacteria are able to utilize the free and bile acid-conjugated taurine as sulfur sources using a plethora of desulfonating enzymes including arylsulfatases, alkylsulfatases, _α_-ketoglutarate-dependent dioxygenases and FMNH_2_-dependent monooxygenases (32, 33). While these enzymes are conserved in fungi, their contribution to sulfonate utilization specifically in pathogenic fungi remains unexplored (34, 35). The only investigation providing genetic evidence of the contribution of fungal desulfonating enzyme was made in *S. cerevisiae* and underscored the role of the _α_-ketoglutarate-dependent dioxygenase Jlp1 in sulfonates utilization (17).

Although considerable advances were made on the metabolic pathways that mediate the utilization of alternative sulfur sources in bacteria, many aspects of this metabolism remain understudied in fungi. To explore the genetic circuit by which *C. albicans* controls sulfur utilization, we characterized the transcriptional atlas associated with sulfur starvation in this yeast. We identified Met32 as a master transcriptional regulator that governs the utilization of different sulfur sources found in different niches of the human host. Importantly, we found that the role of Met32 in the control of sulfur utilization metabolism has diverged from its known function as a transcriptional regulator of *MET* genes in *S. cerevisiae*. This work identified the first major facilitator superfamily member (MFS) transporter in fungi that uptakes GSH under the control of Met32. We also uncovered different desulfonation enzymes that were differentially transcribed under sulfur starvation, and showed that Jlp12, an _α_-ketoglutarate dioxygenase and a target of Met32, was critical for the utilization of alternative sulfur sources. Furthermore, we found that Met32 modulates *C. albicans’* tolerance to reactive sulfur species (RSS) by tuning the superoxide dismutase Sod1 basal transcriptional level. Overall, our study delineates a new mechanism by which fungal pathogens utilize sulfur sources and underscores its importance for fungal fitness *in vivo*.

## Results

### Genome-wide transcriptional surveys of sulfur starvation and utilization in *C. albicans*

Under sulfur starvation, many fungi upregulate transcripts of sulfur utilization together with *MET* genes to guarantee a sufficient supply of this essential nutrient (34, 36–38). To capture elements of the sulfur utilization pathway in *C. albicans*, we performed RNA-seq profiling in cells growing in the absence or presence of different sulfur sources. Cells growing exponentially in the MDM (minimal dextrose medium) were transferred to grow for one hour in either a sulfur lacking medium (SLD), or in SLD supplemented with different sulfur components including ammonium sulfate ((NH_4_)_2_SO_4_), taurine and equimolar mixture of Methionine and Cysteine (Met/Cys). When comparing the transcriptome of cells under sulfur starving conditions to those growing in sulfur-replete media, a total of 922 transcripts were differentially modulated suggesting a drastic remodeling of *C. albicans* cellular processes to accommodate sulfur scarcity (**Figure 1A-C, Table S1 and S2**). A higher dynamic range of expression levels was observed with Met/Cys and sulfate as compared to taurine (**Figure S1**). Overall, and regardless of the resupplied sulfur source, sulfur starved cells upregulated genes related to different processes of sulfur metabolism, oligopeptide uptake, carbohydrate transport and oxidative stress (**Figure 1D**). Downregulated genes were mainly enriched in functions associated with tRNA modification and both cytoplasmic and mitochondrial protein translation.

**Figure 1.**
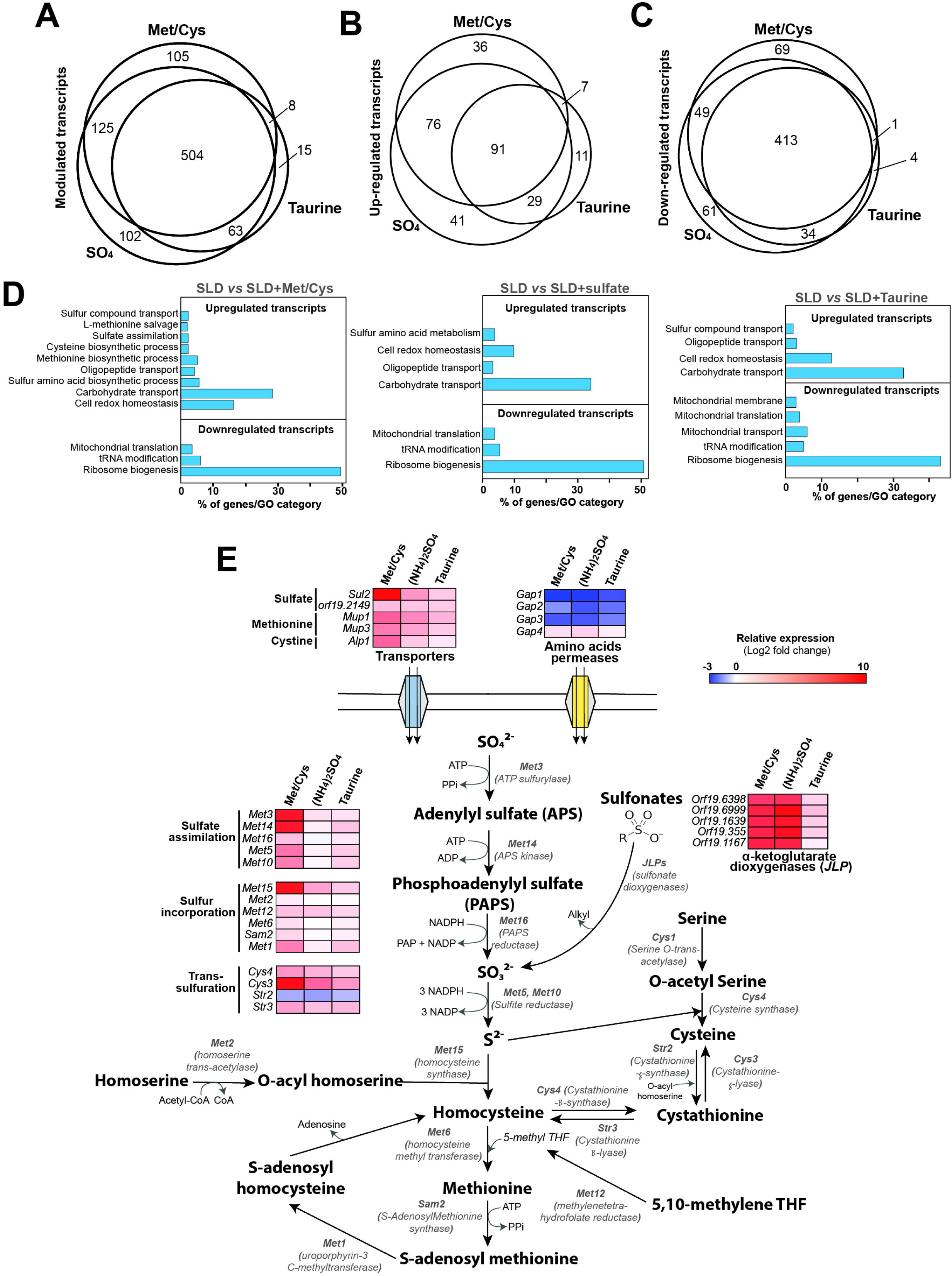
The transcriptional landscape of sulfur starvation in *C. albicans*. (**A-C**) Venn diagram representation of differentially modulated (**A**), upregulated (**B**), and downregulated transcripts (**C**). Differentially expressed transcripts were identified by comparing the transcriptome of *C. albicans* SC5314 cells under sulfur starvation conditions (SLD medium) to that of cells grown in sulfur-replete media (SLD supplemented with either an equimolar mixture of methionine and cysteine (Met/Cys), ammonium sulfate (SO_4_) or taurine) using a false-discovery rate of 1% and a ± 2 log fold-change. (**D**) Gene ontology (GO) analysis of transcripts differentially modulated using the three distinct sulfur sources to mimic sulfur repletion. (**E**) Sulfur utilization and methionine biosynthesis pathways representation. Upregulated (red) and downregulated transcripts (blue) are shown for each sulfur repletion condition. The full list of differentially expressed transcripts is available in **Table S2**.

Globally, under sulfur starvation, *C. albicans* cells are oriented towards sulfur amino acid biosynthesis and the assimilation and uptake of alternative sulfur sources.

#### Sulfur compound uptake and methionine metabolism

Under sulfur deprivation, and when using Met/Cys as sulfur repletion sources, *C. albicans* cells activate genes involved in the different steps of methionine biosynthesis, including sulfur assimilation (*MET3*, *MET14*, *MET16*, *MET5*, *MET10*), incorporation into carbon chain (*MET15*, *MET2, MET6, SAM2, MET1*) and the trans-sulfuration pathway (*CYS4*, *CYS3*) which allow the interconversion between cysteine and homocysteine (**Figure 1E and Table 1).** Different transporters with known roles in sulfur metabolites uptake including the sulfate transporter Sul2, the methionine permeases Mup1 and Mup3 and the cystine transporter Alp1 were significantly upregulated (**Figure 1E)**. Notably, several genes of the methionine salvage pathway that recycles sulfur-containing metabolites to regenerate the cellular methionine pool were also upregulated (*ARO9*, *ADI1*, *MEU1*). A similar transcriptional pattern was observed upon taurine or sulfate repletion. Taken together, these results suggest that under sulfur limitation *C. albicans* cells collectively promote *de novo* synthesis and extracellular uptake of methionine in addition to the recycling of intracellular metabolites to feed the methionine biosynthesis pathway. Homologs of known transcriptional activators of sulfur metabolism in the budding yeast, including the bZIP activator Met4 and the zinc-finger transcription factor Met32, were significantly upregulated when using the three sulfur-replete sources (**Table 1**). Intriguingly, transcript level of *MET30*, a negative regulator of Met4, was also increased under starvation suggesting a time-constrained activity of Met4.

**Table 1.** Summary of sulfur modulated biological functions in *C. albicans.* Black and grey colors underscore genes identified using a Log_2_ FC cutoff > 2 and, <2 but >1, respectively.

|  |  | Met/Cys | Sulfate | Taurine |
| --- | --- | --- | --- | --- |
| **Sulfur metabolic processes** |  |  |  |  |
| **Sulfate transport** |  | *SUL2*, orf19.2149 | *SUL2*, orf19.2149 | *SUL2*, orf19.2149 |
| **Methionine biosynthesis and salvage** | **Sulfate assimilation** | *MET3*, *MET14*, *MET5*, *MET10*, *MET16* | *MET3* | *MET3*, *MET14*, *MET5*, *MET10* |
|  | **Transsulfuration** | *CYS3*, *CYS4* | *CYS3*, *CYS4* | *CYS3*, *CYS4* |
|  | **Sulfate incorporation** | *MET15*, *MET12*, *MET6*, *MET1* | *MET15*, *MET12*, *MET1* | *MET15*, *MET12*, *MET1* |
|  | **tRNA-Methionine** | *tM*(CAU)2 | *tM*(CAU)2 | *tM*(CAU)2 |
|  | **Transcriptional control** | *MET4*, *MET32*, *MET30* | *MET4*, *MET32*, *MET30* | *MET4*, *MET32* |
|  | **Methionine and SAM transport** | *MUP1*, *MUP3* | *MUP1*, *MUP3* | *MUP1*, *MUP3* |
|  | **Methionine salvage** | *ARO9*, orf19.1180, *ADI1*, *MEU1* | *ARO9*, orf19.1180, *ADI1*, *MEU1* | *ARO9*, orf19.1180, *ADI1*, *MEU1* |
| **Sulfonate utilization** | **α-ketoglutarate dioxygenase** | orf19.6999 (*JLP22*), orf19.1167 (*JLP11*), orf19.355 (*JLP21*), orf19.1639 (*JLP13*), orf19.6398 (*JLP12*) | orf19.6999 (*JLP22*), orf19.1167 (*JLP11*), orf19.355 (*JLP21*), orf19.1639 (*JLP13*), orf19.6398 (*JLP12*) | orf19.6999 (*JLP22*), orf19.355 (*JLP21*), orf19.1639 (*JLP13*), orf19.6398 (*JLP12*) |
|  | **Arylsulfatase** | *AYS1* (orf19.1608) | *AYS1* (orf19.1608) | *AYS1* (orf19.1608) |
|  | **Benzene desulfurase** | *SOX1* (orf19.1365), *SOX2* (orf19.3901) | *SOX1* (orf19.1365), *SOX2* (orf19.3901) | *SOX1* (orf19.1365), *SOX2* (orf19.3901) |
|  | **Flavin-containing monooxygenase** | orf19.2197 (*FMO3*), orf19.7364 (*FMO5*), *SSP96* | *FMO1*, orf19.2197 (*FMO3*), orf19.7364 (*FMO5*), *SSP96* | *FMO1*, *FMO2*, orf19.2197 (*FMO3*), orf19.7364 (*FMO5*), *SSP96* |
| **Glutathione** | **Synthesis** | *GCS1*, *GSH2* | *GCS1*, *GSH2* | *GSH2* |
|  | **Transferase** | *GTT1*, *GST2*, *GTT13*, *GST3* | *GTT1*, *GST2*, *GTT13*, *GST3* | *GST2*, *GTT13*, *GST3* |
| **Disulfide reductases** | **Glutaredoxin** | *GRX1* | *GRX1* | *GRX1* |
|  | **Thioredoxin** | *TRP99* | *TRP99* | - |
| **Cysteine hyperoxidation** | **Sulfiredoxin** | *SRX1* (orf19.3537) | *SRX1* (orf19.3537) | *SRX1* (orf19.3537) |
|  | **Peroxiredoxin** | *AHP2* | *AHP2* | *AHP2* |
| **Methionine sulfoxide reductase** | **Methionine-S-sulfoxide reductase** | *MXR1* | - | - |
|  | **Methionine-R-sulfoxide reductase** | orf19.5553, *MXR2* (orf19.3292) | orf19.5553, *MXR2* (orf19.3292) | orf19.5553 |
| **Nitrogen and peptide metabolic processes** |  |  |  |  |
| **Nitrogen metabolism** | **Peptide transporters** | *OPT1*, *OPT2*, *OPT3*, *OPT4*, *OPT5*, *OPT7*, *OPT8*, *OPT9*, *PTR2* | *OPT1*, *OPT2*, *OPT3*, *OPT4*, *OPT5*, *OPT7*, *OPT8*, *OPT9*, *PTR2* | *OPT1*, *OPT2*, *OPT4*, *OPT5*, *OPT7*, *OPT8*, *OPT9*, *PTR2* |
|  | **Allantoate transporters** | *SEO1*, *SEO2*, *SOA1*, *DAL5*, orf19.7158, *DAL9*, orf19.6522, *SEO13* | *SEO1*, *SEO2*, *SOA1*, *DAL5*, orf19.7158, *DAL9*, orf19.6522, *SEO13* | *SEO2*, *SOA1*, *DAL5*, orf19.7158, *DAL9*, orf19.6522, *SEO13* |
|  | **Allantoate degradation** | *DAL2*, *DAL3* | *DAL2* | *DAL2* |
|  | **transporters** | *DUR35, DUR4* | *DUR35, DUR4* | *DUR35, DUR4* |

**Table 2.** Met32 DNA-bound promoters associated with sulfur metabolism identified by ChEC-seq. Transcript levels of the RNA-seq experiments are indicated both in the WT and the *met32* mutant. **^a^** Transcript levels from the *met32* vs WT (SLD + taurine) comparison.

| Gene ID | Gene name | Function | CheC binding | RNA-seq expression |  |  |  |
| --- | --- | --- | --- | --- | --- | --- | --- |
|  |  |  |  | Met/Cys | SO<sub>4</sub> | Taurine | *met32*<sup>a</sup> |
| CR_08340W_A | CYS3 | Cystathionine gamma-lyase | 4246.6 | 5.19 | 2.97 | 1.98 | 4.12 |
| CR_09170C_A | SSU1 | Protein similar to *S. cerevisiae* Ssu1 sulfite efflux pump | 1296.5 | 0.70 | -0.02 | -0.43 | -0.70 |
| C1_07120W_A | GAP4 | High-affinity S-adenosylmethionine permease | 3232.9 | 0.91 | 1.29 | 0.56 | -0.65 |
| C1_11870W_A | MUP1 | High affinity methionine permease | 3010.5 | 3.99 | 3.02 | 1.93 | 0.30 |
| C4_00440C_A | OPT7 | Putative oligopeptide transporter; possibly transports GSH or related compounds | 1945.9 | 6.95 | 4.24 | 1.51 | 1.73 |
| C7_02000C_A | SOA1 | Putative allantoate permease | 929 | 12.78 | 13.12 | 2.11 | -2.02 |
| CR_08310C_A | JLP12 | Predicted sulfonate α-ketoglutarate dioxygenase | 3106.1 | 9.50 | 10.02 | 2.30 | -4.30 |
| C1_11530C_A | JLP11 | Predicted sulfonate α-ketoglutarate dioxygenase | 1293.4 | 10.97 | 11.11 | 0.95 | -5.19 |
| C3_02360C_A | AYS1 | Arylsulfatase | 1225.7 | 9.86 | 8.90 | 2.57 | 4.95 |
| C3_05840W_A | FMO5 | Protein with predicted N,N-dimethylaniline monooxygenase activity | 1225.7 | 10.78 | 11.51 | 3.61 | -9.51 |
| C1_13870W_A | MET3 | ATP sulfurylase; involved in sulfate assimilation | 1803.7 | 8.10 | 1.14 | 1.03 | 4.93 |
| C4_04000W_A | MET4 | Putative transcription co-activator; predicted role in sulfur amino acid metabolism | 2093.3 | 3.23 | 1.95 | 1.84 | 2.19 |
| C2_10230W_B | MET32 | General Zinc-finger DNA-binding transcription factor | 1726.9 | 5.54 | 5.70 | 2.31 | -14.32 |
| C1_01130W_A | MET30 | Putative ubiquitin ligase complex component | 994.5 | 2.44 | 2.11 | 0.86 | 0.33 |
| C4_02990C_B | GST2 | Glutathione S transferase | 2066 | 5.94 | 5.27 | 1.99 | -2.05 |
| CR_08370W_A | GSH2 | Putative glutathione synthase | 993.7 | 2.34 | 2.21 | 1.78 | 0.38 |
| C2_05060C_A | SRX1 | Putative sulfiredoxin | 3016.1 | 3.29 | 2.98 | 1.77 | 1.15 |
| C7_02810W_A | PRX1 | Thioredoxin peroxidase | 1128.6 | 1.64 | 0.63 | 0.01 | -0.68 |
| C7_02390W_A | AHP2 | Putative thiol-specific peroxiredoxin | 1077.6 | 2.95 | 2.70 | 1.51 | -1.90 |
| C1_10800C_A | ALP1 | Cystine transporter | 1007.8 | 3.95 | 1.44 | 0.73 | -0.58 |
| CR_09010C_A |  | Putative cystathionine gamma-synthase | 994.7 | 1.99 | 1.64 | 0.84 | 0.90 |
| CR_04500C_A |  | 2-OG-Fe(II) oxygenase | 2277.7 | 5.02 | 3.96 | 1.51 | 2.40 |
| CR_06660W_A | SEO1 | Protein with similarity to permeases | 952.2 | 2.90 | 2.17 | 0.77 | 1.68 |
| CR_10770W_A | SEO13 | Predicted MFS membrane transporter | 1419.1 | 8.24 | 7.83 | 2.68 | -2.64 |
| C4_02320C_B | SOD1 | Cytosolic copper- and zinc-containing superoxide dismutase | 3181.8 | 1.27 | 1.39 | 0.44 | -0.45 |
| CR_08320W_B | ATS1 | Protein required for modification of wobble nucleosides in tRNA | 2764.4 | 1.47 | 1.68 | 2.01 | 0.97 |

#### Sulfur utilization genes

Sulfur deprivation elicited upregulation of many transcripts involved in the utilization of alternative sulfur sources. Transcriptional activation of homologs of the *S. cerevisiae* Jlp1, a Fe(II)-dependent sulfonate/_α_-ketoglutarate dioxygenase, was observed under all the three sulfur-replete sources. *C. albicans* genome encodes five Jlp1 homologs which were significantly upregulated in our study (these genes were named as following: *JLP11, JLP12, JLP13, JLP21, JLP22*; **Table 1 and Figure S2**). Interestingly, the ortholog of the recently identified sulfonate transporter Soa1 in *S. cerevisiae* (orf19.6520), was highly upregulated. Genes encoding other homologs of bacterial enzymes involved in sulfur release from organic compounds including the arylsulfatase *AYS1*, two homologs of the bacterial alkanesulfonate monooxygenases (*SOX1* and *SOX2*), and five FMNH_2_-dependent monooxygenases (*FMO1*, *FMO2*, *FMO3*, *FMO5*, *SSP96*) were also upregulated.

#### Nitrogen metabolism

Different biological functions associated with nitrogen utilization processes were also overrepresented in our datasets under sulfur limitation, reflecting a coordinated control of nitrogen and sulfur metabolism. The transcriptional response under the three sulfur-replete conditions reflects a derepression of the *bona fide* nitrogen catabolite repression (NCR) response, a situation allowing fungal cells to indiscriminately scavenge alternative, nonpreferred nitrogen sources to survive (18). Transcript levels of many transporters of peptides, urea, and allantoate were highly activated along with genes involved in nitrogen sources utilization (**Table 1**).

#### Redox processes

Transcripts associated with the maintenance of the redox homeostasis included mainly genes of GSH biosynthesis (*GCS1*, *GSH2*) and reduction of GSH disulfide (*GRX1*, *TRP99*) in addition to glutathione S-transferases (*GST2, GST3, GTT1, GTT13*) (**Table 1**). Activation of genes involved in both GSH biosynthesis and the recycling of its oxidized form (GSSG) might reflect a cellular compensatory response as sulfur depletion led to a shortage of cysteine, one of the three amino acids components of GSH. This is also supported by the fact that *OPT7* required for GSH uptake in *C. albicans* was among the highly upregulated transcripts (39). Sulfur starvation induced many genes known for their role in protecting protein sulfur residues against oxidation by reactive oxygen species including methionine sulfoxide reductases (*MXR1*, *MXR2*, orf19.5553) and proteins of the sulfiredoxin-peroxiredoxin redox system.

### Transcriptomic-guided discovery of sulfur utilization genes

As our transcriptional profiling underlined many putative fungal desulfonation enzymes orthologous to known bacterial organosulfonates utilization proteins, we decided to test their contribution to sulfonate utilization in *C. albicans*. Mutants of Fe(II)-dependent sulfonate/_α_-ketoglutarate dioxygenases (*jlp11*, *jlp12*, *jlp13, jlp21, jlp22*), alkanesulfonate monooxygenases (*sox1*, *sox2*), FMNH_2_-dependent monooxygenases (*fmo1*, *fmo2*, *fmo3*, *fmo5*, *ssp96*) and the arylsulfatase (*ays1*), were tested for taurine utilization. As all these enzymes are monooxygenases that catalyze oxygen-dependent desulfonation, we tested their growth under both normoxia and hypoxia (5% O_2_), the prevalent environmental condition that *C. albicans* encounters in most of the colonized niches. Our data show that, except for *jlp12* that was unable to utilize taurine as sole sulfur source, all other tested mutants had growth patterns comparable to that of the WT strain (**Figure 2A**). Interestingly, failure of *jlp12* to utilize taurine was perceived exclusively under hypoxia suggesting an important role of this enzyme for *C. albicans* fitness inside the host. In addition to taurine, *jlp12* failed to utilize other sulfonates including benzene sulfonate and taurocholate (**Figure 2B**). Furthermore, *jlp12* demonstrated the capacity to assimilate sulfate ((NH_4_)_2_SO_4_), thereby supporting the desulfonation activity of this enzyme (**Figure 2B**). When either methionine or cysteine was supplemented to SLD medium, *jlp12* grew comparably as in the complete MDM medium which further indicates the inability of this mutant to utilize alternative sulfur sources for synthesizing these essential amino acids. The fact that among all differentially transcribed desulfonation enzymes, only Jlp12 is essential for sulfonate utilization might reflect either a functional redundancy or a narrow specificity of the other enzymes toward other sulfur sources yet to be determined.

**Figure 2.**
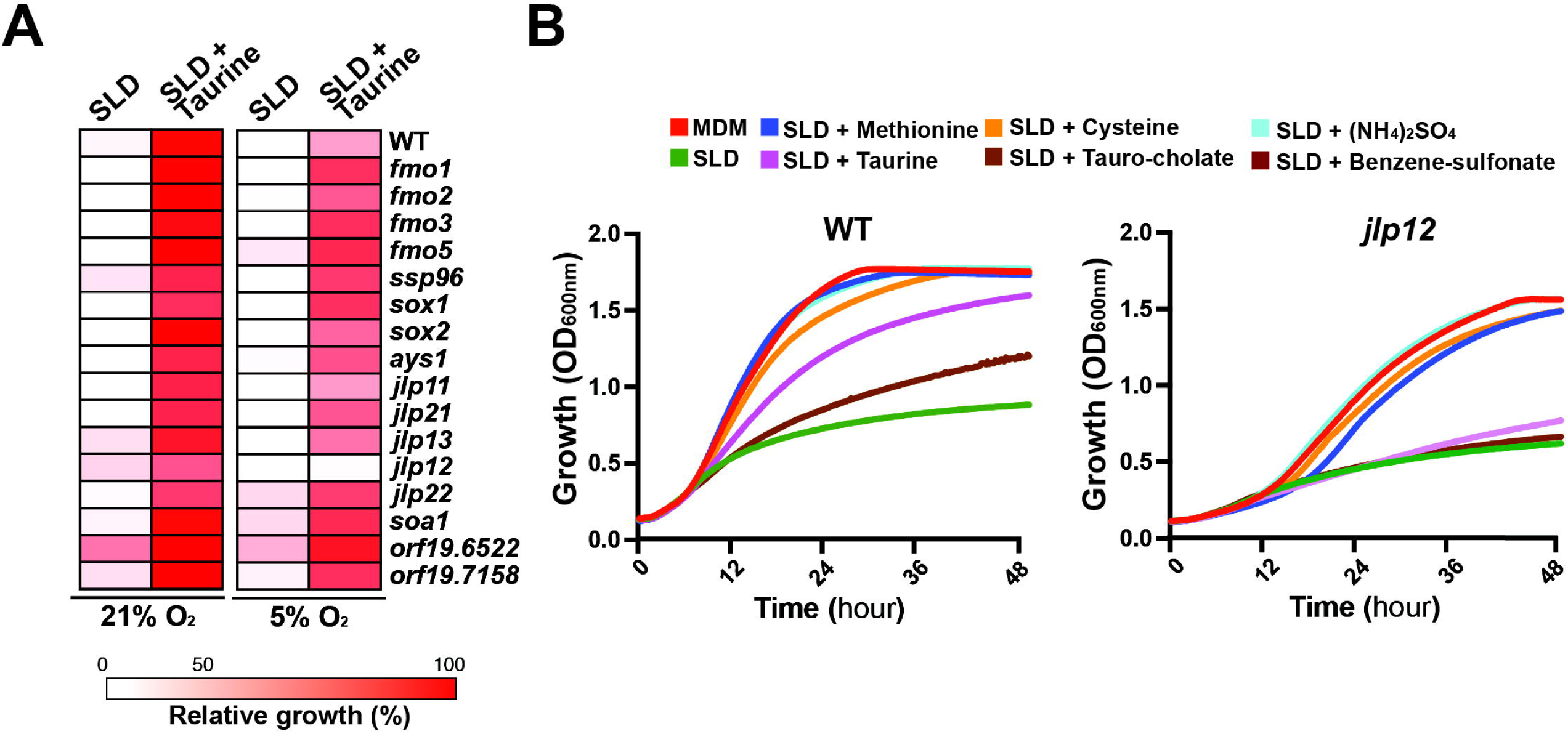
*C. albicans* sulfonate/_α_-ketoglutarate dioxygenase Jlp12 is essential for alternative sulfur utilization. (**A**) Taurine utilization assay of putative desulfonation enzymes that are transcriptionally modulated under sulfur starvation. The heatmap represents relative growth of each mutant as compared to the WT (SN250) strain under both normoxic and hypoxic conditions. Each strain was grown in SLD alone or SLD supplemented with taurine and OD_600_ readings were taken over 48 hours of growth at 30°C. For mutants of the GRACE conditional collection (*fmo1*, *fmo2*, *fmo3*, *fmo5*, *sox1*, *sox2*, *ays1*, *jlp11*, *jlp13*, *jlp21*, *jlp22*, *orf19.6522* and *orf19.7158*), 20 µg/ml doxycycline was added to the culture medium for transcriptional repression. (**B**) *jlp12* mutant is unable to utilize different alternative sulfur sources. Both WT (SN100) and *jlp12* strains were grown in MDM, SLD alone or supplemented with 80 µM of different sulfur sources (taurine, cysteine, methionine, ammonium sulfate, benzene sulfonate and tauro-cholate). Cells were grown at 30°C, and OD_600_ readings were taken every 10 min for 48 hours.

We also examined selected homozygous deletion strains of *C. albicans* orthologs of the recently characterized sulfonate transporter Soa1 in *S. cerevisiae* (16) that were significantly modulated in our RNA-seq experiments. Inactivation of the *C. albicans* Soa1 (orf19.6520) and two other homologous transporters (orf19.6522 and orf19.7158) had no impact on taurine utilization (**Figure 2A**). This finding suggests that either the function of these transporters has diverged in *C. albicans,* or they might be functionally redundant.

### Met32 is required for sulfur utilization in *C. albicans*

Promoter sequences of the top 100 upregulated genes under sulfur starvation were searched for potential cis-regulatory motifs using MEME algorithm (40). Across the three RNA-seq datasets, one particular motif AAYTGTGGCT was statistically overrepresented and corresponds to the motif recognized by the *S. cerevisiae* transcription factor Met32, a master regulator of *MET* genes (19) (**Figure 3A and Table S3**). Previous work on *C. albicans* had excluded the contribution of Met32 as a transcriptional regulator of methionine biosynthetic genes, as genetic inactivation of *MET32* did not result in methionine auxotrophy (26). We found that the Met32 motif was present in the promoters of only three *MET* genes (*MET15*, *MET10, MET14*) suggesting a minor contribution to *MET* gene regulation as compared to *S. cerevisiae* (41). We confirmed that *met32* did not exhibit methionine auxotrophy and grew comparably to the WT parental strain when either methionine or both methionine and cysteine were omitted from the growth medium (**Figure 3B**). In contrast, no growth was observed for *cbf1* mutant, which is known to have methionine auxotrophy and served as the control (28). We hypothesized that Met32 might govern transcriptional control of sulfur utilization, and its function has diverged from its known role as a transcriptional regulator of *MET* genes in *S. cerevisiae*. *met32* cells were unable to utilize either free or bile acid-conjugated taurine (taurocholic acid and taurodeoxycholic acid) in addition to sodium benzenesulfonate and the sulfoxide DMSO, and exhibited a growth pattern comparable to cells growing in the sulfur free medium, SLD (**Figure 3C**).

**Figure 3.**
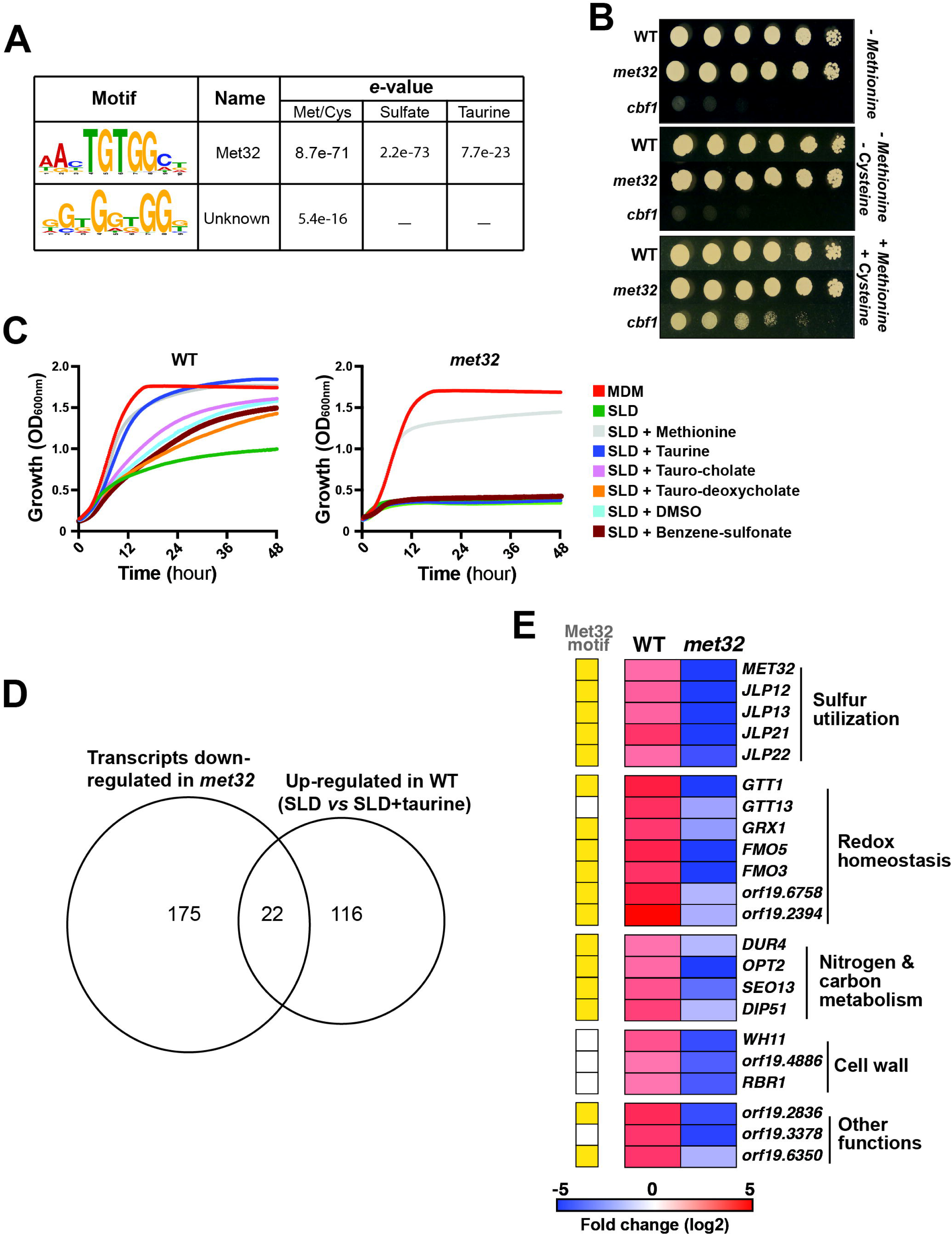
Met32 governs the transcriptional control of sulfur utilization metabolism in *C. albicans*. (**A**) *De novo* prediction of DNA cis-regulatory motifs enriched in the top 100 highly upregulated transcripts under sulfur starvation using MEME analysis tool (http://meme-suite.org). The two significant sequence logos identified by MEME are shown together with their respective e-values. (**B**) Methionine auxotrophy assay for *met32* and *cbf1* mutants. WT and mutant strains were serially diluted and spotted on SD-agar medium lacking either methionine or both methionine and cysteine and incubated for 24 hours. (**C**) utilization of various alternative sulfur sources by *met32*. WT (SN250) and *met32* strains were grown in MDM, SLD alone or supplemented with 80 µM of different sulfur sources. Cells were incubated at 30°C, and OD_600_ readings were taken every 10 minutes for 48 hours. (**D**) Met32-dependent modulation of transcripts upregulated under sulfur starvation. Venn diagram showing overlaps between upregulated sulfur responsive transcripts in the WT strain (**Table S3**) and those downregulated in *met32* under sulfur starvation (SLD) as compared to taurine-repleted condition (**Table S2**). Modulated transcripts in *met32* mutant were identified using a false-discovery rate of 5% and a ± 1.5 log fold-change cutoff. (**E**) Heatmap representation of Met32-dependent transcripts and their biological functions. Presence of the Met32-predicted cis-regulatory motif is indicated in yellow.

To further investigate the role of Met32 in the transcriptional control of sulfur utilization, we performed transcriptional profiling of both WT and *met32* cells growing in SLD medium supplemented with taurine. We found that Met32 was required to activate and repress 197 and 283 transcripts, respectively (**Table S3**). Among genes that *met32* failed to activate, a total of 22 were upregulated upon taurine utilization by the WT strain (**Figure 3D and Table S3**). This subset of transcripts was enriched in gene encoding _α_-ketoglutarate dioxygenases (*JLP12, JLP13*, *JLP21*, *JLP22*), the flavin-containing monooxygenases (*FMO5*, *FMO3*), and genes functioning in redox homeostasis such as the glutathione S-transferases (*GTT1*) and the glutaredoxin *GRX1* (**Figure 3E**). As *JLP12* was essential for taurine utilization (**Figure 2B**), our data suggest that Met32 might govern the transcriptional control of sulfonate utilization by acting as an activator of *JLP12* under sulfur starvation.

### Met32 governs transcriptional control of sulfur utilization in *C. albicans*

To investigate the direct regulation of sulfur utilization genes by Met32, we performed ChEC-seq (Chromatin endogenous cleavage sequencing) analysis to identify promoters of genes occupied by this transcription factor. Promoters bound by Met32 were enriched for genes involved in sulfur metabolism, cell redox defense and biofilm formation (**Figure 4A and Table S4**). Met32 binds the promoters of different desulfonating enzymes, including the sulfonate/_α_-ketoglutarate dioxygenases (Jlp11 and Jlp12), the FMNH2-dependent monooxygenase Fmo5 and the arylsulfatase Ays1 (**Figure 4B-C and Table S4**). Met32 was also found in the promoters of genes related to glutathione and methionine metabolism, transport of sulfur-containing metabolites and antioxidative defense (**Figure 4B**). Interestingly, Met32 occupies its own promoter and the promoters of the transcription factor Met4 and the ubiquitin ligase Met30 (**Figure 4C)**. To uncover the DNA-binding motif of Met32, the 120 most enriched peaks were analyzed for motif enrichment using MEME (40). We found that Met32 binding sites detected by ChEC were enriched for the motif RBTGTGGYKB which resembles that of Met32 of *S. cerevisiae* (19) (**Figure 4D**) and to that enriched in the promoter of upregulated transcripts in response to sulfur starvation (**Figure 3A**). Of note, we found that Met32 occupied the promoter regions of 10 sulfur-related genes that were activated upon sulfur starvation when using taurine as a repletion sulfur source (**Figure 4E**). Together, these indicate that Met32 directly modulates sulfur utilization genes through a conserved DNA recognition motif.

**Figure 4.**
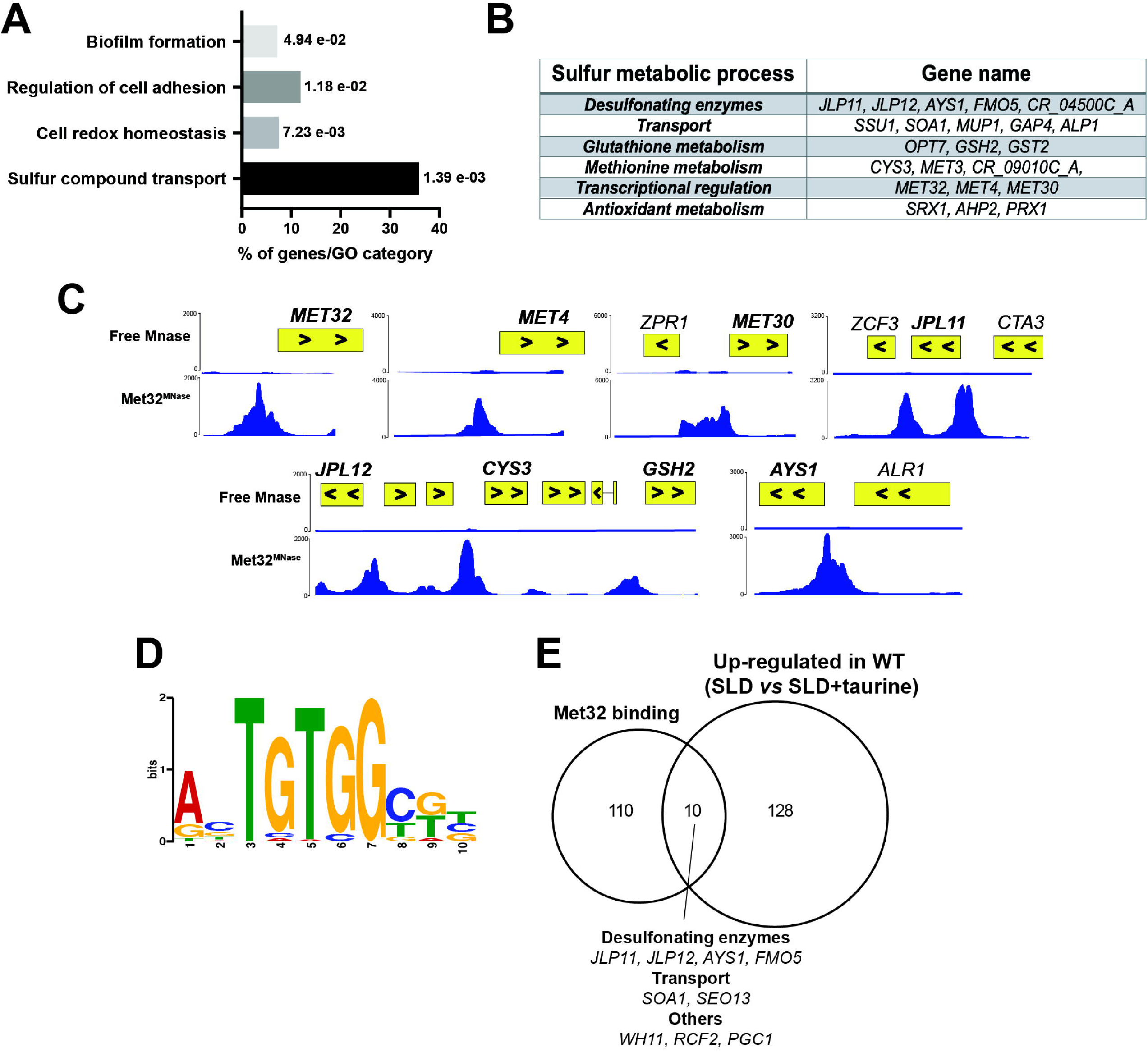
Genome-wide occupancy of Met32 under sulfur starvation. **(A)** Gene ontology analysis of gene promoters bound by Met32. Met32 binding was assessed by ChEC-seq in Met32-MNase-tagged strain growing in SLD medium at 30°C. (**B**) List of Met32 target genes associated with sulfur metabolic processes. (**C**) Snapshots of genomic regions showing Met32 binding to intergenic regions of genes involved in sulfur metabolism. ChEC-seq signals are shown for the Met32^MNase^ and the control stain (free MNase). (**D**) Motif enriched in the promoters of Met32 binding loci. The motif logos were generated using MEME-ChIP software on the top 120 high-scoring peaks. (**E**) Venn diagram showing the overlaps between Met32-bound promoters and upregulated sulfur responsive transcripts in WT under sulfur starvation (SLD) when compared to taurine-repleted conditions (SLD + Taurine).

### *C. albicans* utilization and uptake of glutathione requires Met32

The thiol tripeptide GSH is a highly abundant metabolite and sulfur source that is found at millimolar levels in many anatomical sites of the human host (42). As Met32 occupied the promoter of Opt7 (**Figure 4B**), a GSH transporter in *C. albicans* (39), we examined the ability of the *met32* mutant to utilize glutathione (GSH) as a sole source of sulfur. Consistent with prior works (43), we found that *C. albicans* utilizes GSH as a sole sulfur source (**Figure 5A**). Conversely, the *met32* mutant was unable to thrive in SLD supplemented with GSH suggesting that Met32 is required for extracellular GSH utilization (**Figure 5A**). This finding suggests that GSH uptake and/or degradation processes are impaired in *met32* mutant. In SLD medium, we found that *met32* accumulated twice less GSH as compared to the WT strain (**Figure 5B**). When GSH was added to the SLD medium, the intracellular GSH content significantly increased in the WT, whereas the *met32* mutant showed similar GSH levels to those observed in the SLD medium alone (**Figure 5B)**. Thus, Met32 appears to be required for GSH utilization through the modulation of its uptake.

**Figure 5.**
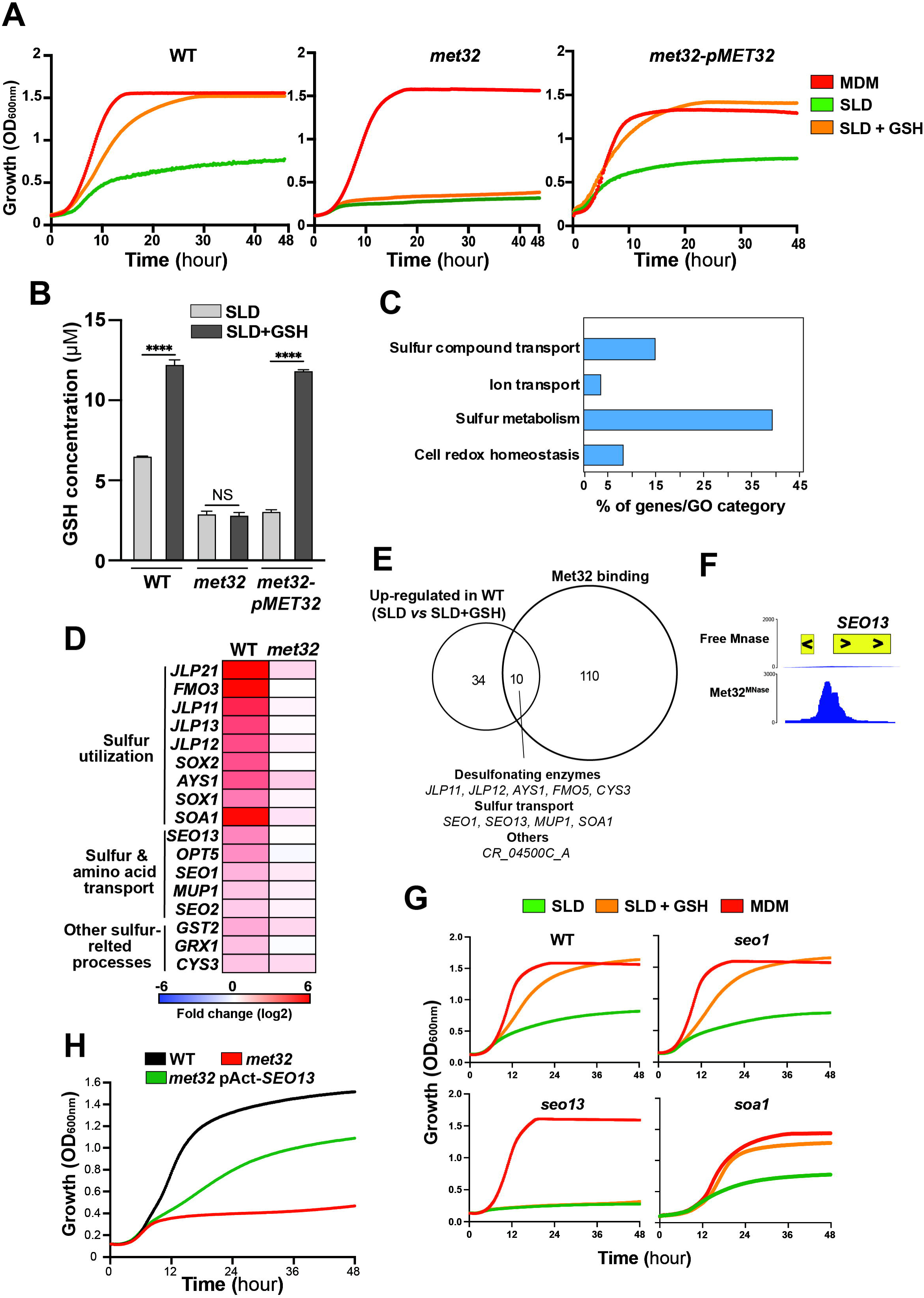
Met32 controls the uptake of glutathione through the MFS transporter Seo13. (**A**) *met32* is unable to utilize GSH as a sole source of sulfur. WT (SN250), *met32* and *met32*-p*MET32* complemented strains were grown in MDM, SLD alone or SLD supplemented with 80 µM GSH at 30°C. OD_600_ readings were recorded every 10 min for 48 hours. (**B**) Met32 modulates GSH content in *C. albicans*. GSH levels were measured in WT (SN250), *met32* and *met32*-p*MET32* complemented strains grown in either SLD or SLD + 80 µM GSH. Data represent the mean ± standard deviation of at least three independent biological replicates. Statistical analysis was performed using one-way ANOVA test with **** P-value < 0.00001; NS: nonsignificant. (**C**) Gene ontology (GO) analysis of differentially modulated transcripts in WT (SN250) cells grown under sulfur starvation (SLD medium) versus GSH-replete conditions (SLD+GSH). Differentially expressed genes were identified using a false-discovery rate of 5% and a log_2_ fold-change cutoff of ±1. The full list of differentially expressed transcripts is available in **Table S5**. (**D**) Heatmap representation of sulfur metabolism-related transcripts requiring Met32 for their activation. (**E**) Venn diagram showing the overlap between Met32-bound promoters and upregulated sulfur responsive transcripts under sulfur starvation (SLD) compared to GSH-repleted condition (SLD + GSH). (**F**) Binding of Met32 to the intergenic region of the MFS transporter Seo13. (**G**) Mutant of the MFS transporter Seo13 fails to utilize GSH as a sole sulfur source. WT (CAI4) and GRACE conditional mutants (*seo13*, *soa1*, *seo1*) were grown in MDM, SLD alone or supplemented with 80 µM GSH, and OD_600_ readings were taken every 10 min for 48 hours. 20 µg/ml doxycycline was added to repress gene expression (77). (**H**) Increasing *SEO13* dosage in *met32* restore partially the GSH utilization defect. WT (WT-Cip-Act), *met32* and *met32* overexpressing *SEO13* (*met32*-pAct-*SEO13*) were grown in SLD supplemented with 80 µM GSH at 30°C. OD_600_ readings were taken every 10 min for 48 hours.

### Met32 controls GSH uptake through the MFS transporter Seo13

As Opt7 is the only characterized GSH transporter in *C. albicans*, we hypothesized that Met32’s requirement for GSH uptake could be mediated by direct modulation of *OPT7* expression. First, we assessed the transcript levels of *OPT7* using qPCR to test whether the failure of *met32* to grow on medium with GSH as a sole sulfur source is related to inappropriate regulation of this gene. Expression levels of *OPT7* were not impacted in *met32* as compared to the WT strain in either SLD or GSH-supplemented SLD (**Figure S3A**). Second, we tested whether constitutive expression of *OPT7* might rescue the ability of *met32* mutant to utilize GSH. We found that overexpressing *OPT7* using the actin promoter in *met32* did not rescue the GSH utilization defect of this mutant (**Figure S3B**). Together, this suggests that Met32 is unlikely modulating GSH uptake through Opt7.

To gain a mechanistic insight into the role of Met32 in mediating GSH uptake, we performed an RNA-seq transcriptional profiling in both WT and *met32* mutant strains growing in SLD medium supplemented or not with GSH. First, we underlined the transcriptional profile of GSH utilization in *C. albicans,* by comparing the transcriptome of WT cells in SLD to that of cells growing in GSH-repleted medium. Compared to taurine, Cys/Met, and ammonium sulfate, only a few transcripts were modulated when using GSH as a sole source of sulfur (44 upregulated genes; **Table S5**). Upregulated transcripts were enriched mainly in sulfur metabolism, cell redox homeostasis and transport (**Figure 5C**). Induction of the majority of the responsive genes was altered in *met32* specifically transcripts related to sulfur metabolism and transport (**Figure 5D and Figure S3C**). Among the GSH-responsive genes that were upregulated, 10 genes were bound by Met32 (**Figure 5E**). This includes the desulfonating enzymes (*JLP11*, *JLP12*, *FMO5*, *AYS1*), the methionine transporter *MUP1*, the MFS transporters *SEO1* and *SEO13,* the cystathionine gamma-lyase *CYS3,* the sulfonate transporter *SOA1* and the uncharacterized gene CR_04500C_A (**Figure 5E-F**). Among the transporters that rely on Met32 for their activation and that are direct targets of this transcription factor, we tested the mutants of the two uncharacterized MFS transporters *Seo1* and *Seo13*, as well as the recently characterized sulfonate transporter Soa1 in *S. cerevisiae* (16), for their ability to thrive on GSH as a sole source of sulfur. We found that *seo1* and *soa1* mutants exhibited a growth pattern similar to their parental WT strains, while *seo13* showed a notable growth defect similar to *met32* (**Figure 5G**). These data indicate that Met32 modulates GSH uptake in *C. albicans* by transcriptionally tuning the MFS transporter *SEO13*. Consistently, increasing the dosage of *SEO13* partially rescued growth defect of *met32* in SLD medium with GSH as a sole sulfur source (**Figure 5H**).

### Met32 mediates sulfite resistance independently from Zcf2-Ssu1 circuit

We found that Met32 occupied the promoter of the sulfite efflux pump Ssu1, which is essential for *C. albicans* to tolerate sulfite and reactive sulfur species (RSS) (44, 45) (**Figure 4B**). Thus, we hypothesized that Met32 might modulate sulfite and RSS tolerance. We found that *met32* exhibited increased sensitivity to sulfite, similar to the *ssu1* mutant, when compared to the WT strain (**Figure 6A**). This suggests that Met32 might modulate sulfite tolerance through the transcriptional regulation of *SSU1*. qPCR analysis revealed that *SSU1* inducibility in response to sulfite excess was not impaired in *met32* as compared to the WT strain, questioning the functional link between Ssu1 and Met32 (**Figure 6B**). Genetic inactivation of the transcription factor *ZCF2*, which governs sulfite tolerance by controlling Ssu1 (44, 45), in *met32* led to a synthetically sick *met32 zcf2* double mutant suggesting that both Met32 and Zcf2-Ssu1 circuit operate in parallel pathways (**Figure 6C**). Together, these data suggest that *met32* hypersensitivity to sulfite is unlikely related to impaired transcriptional control of *SSU1*.

**Figure 6.**
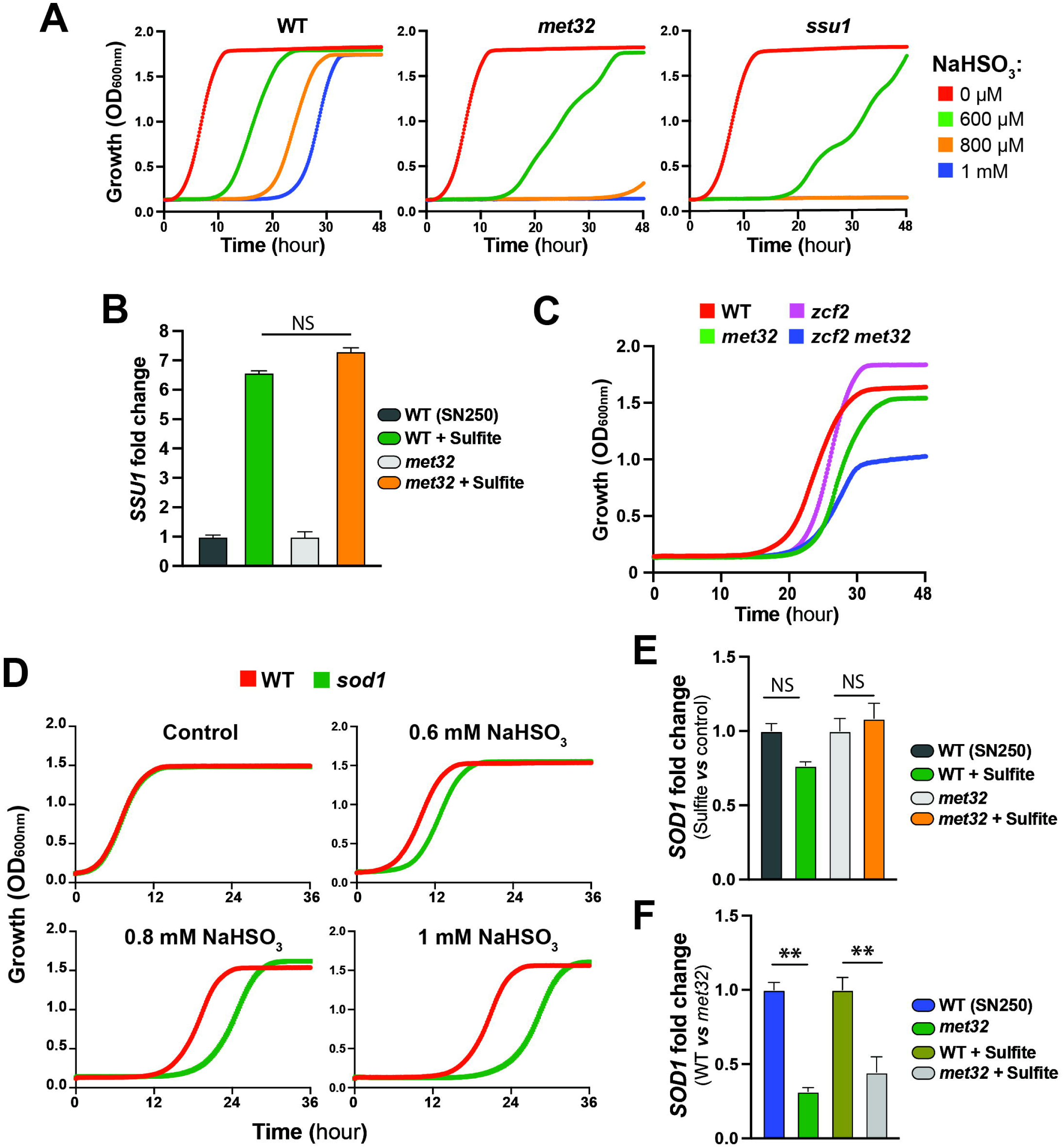
Met32 modulates *C. albicans* tolerance to sulfite. (**A**) *met32* is highly sensitive to sulfite excess. Cultures of the indicated strains were grown in SC medium supplemented with different concentrations of sodium bisulfite (NaHSO_3_) and OD_600_ readings were monitored for 48 hours. (**B**) Transcript levels of the sulfite efflux transporter *SSU1* are not impaired in *met32* mutant. Transcript levels of *SSU1* were measured by qPCR and fold-changes were calculated using the comparative Ct method. For each strain, data were normalized relative to the untreated condition. (**C**) Genetic inactivation of *ZCF2* in *met32* results in a synthetic sick phenotype. WT (SN250), *met32*, *zcf2* and the double mutant *zcf2 met32* were grown in SC medium with 1mM sodium bisulfite. OD_600_ readings were monitored over 48 hours. (**D**) Sod1 is required for resistance to sulfite excess. Strains were grown in SC medium supplemented with different concentrations of sodium bisulfite. (**E-F**) Met32 modulates *SOD1* basal transcription. *SOD1* transcript level was assessed by qPCR. For each strain, data were normalized relative to either the untreated condition (**E**) or to the WT strain (**F**). Data were generated from at least two independent biological replicates and then expressed as means ± standard deviation. Statistics are a one-way ANOVA analysis with ** p < 0.01; NS: nonsignificant.

In *S. cerevisiae*, the superoxide dismutase Sod1 is essential to protect cells from sulfide- and RSS toxicity (46). However, in *C. albicans* such function was not characterized yet. As Met32 bound the promoter of the homologue of *S. cerevisiae* Sod1, we hypothesized that resistance to excess sulfite might rely on the transcriptional control of *SOD1* by Met32. First, we tested the requirement of Sod1 on the ability of *C. albicans* cells to resist elevated concentrations of sulfite. We found that *sod1* mutant exhibited hypersensitivity to sulfite, although to a lesser degree than *met32* mutant (**Figure 6D**). In accordance with previous work (45), our qPCR analysis uncovered that *SOD1* transcript was not modulated by sulfite excess (**Figure 6E**). However, we found that *SOD1* was downregulated in *met32* mutant as compared to WT parental strain, with a similar level of downregulation observed under both treated and untreated conditions (**Figure 6F**). Taken together, these observations suggest that Met32 modulates *C. albicans* tolerance to RSS by controlling *SOD1* basal transcriptional level. Furthermore, this underscores the dual role of the transcription factor Met32 in coordinating both the catabolism of sulfur-containing metabolites and the detoxification of the resulting RSS-derived catabolites.

### Genetic inactivation of *MET32*, *JLP12* and *SEO13* led to attenuated virulence

To elucidate the importance of sulfur utilization processes in the ability of *C. albicans* to infect the host, we tested *met32* virulence in the *Galleria mellonella* model for systemic infection. The *met32* mutant displayed attenuated virulence as compared to WT especially at 1–3-day interval (**Figure 7A**). At day 1 post-infection, the *met32* mutant exhibited a mortality rate of only 12.5% while the WT resulted in 45% mortality. Reintegration of *MET32* in *met32* mutant restored *C. albicans* virulence to a level comparable to the WT strain. We also tested the ability of *met32* mutant to cause damage to human HT-29 enterocytes using LDH release assay. We found that *met32* cells caused slightly, but significantly, less damage to HT-29 enterocytes compared to the WT strain and the complemented strain (**Figure 7B**). Interestingly, when methionine was added to the pre-cultures, *met32* exhibited an invasion capacity similar to that of the WT strain. This suggests that *met32* invasion defect is most likely due to its inability to acquire sulfur from substrates for the synthesis of the proteogenic amino acid methionine. Together, these results suggest that sulfur utilization metabolism is important for disseminated infection and host cell invasions.

**Figure 7.**
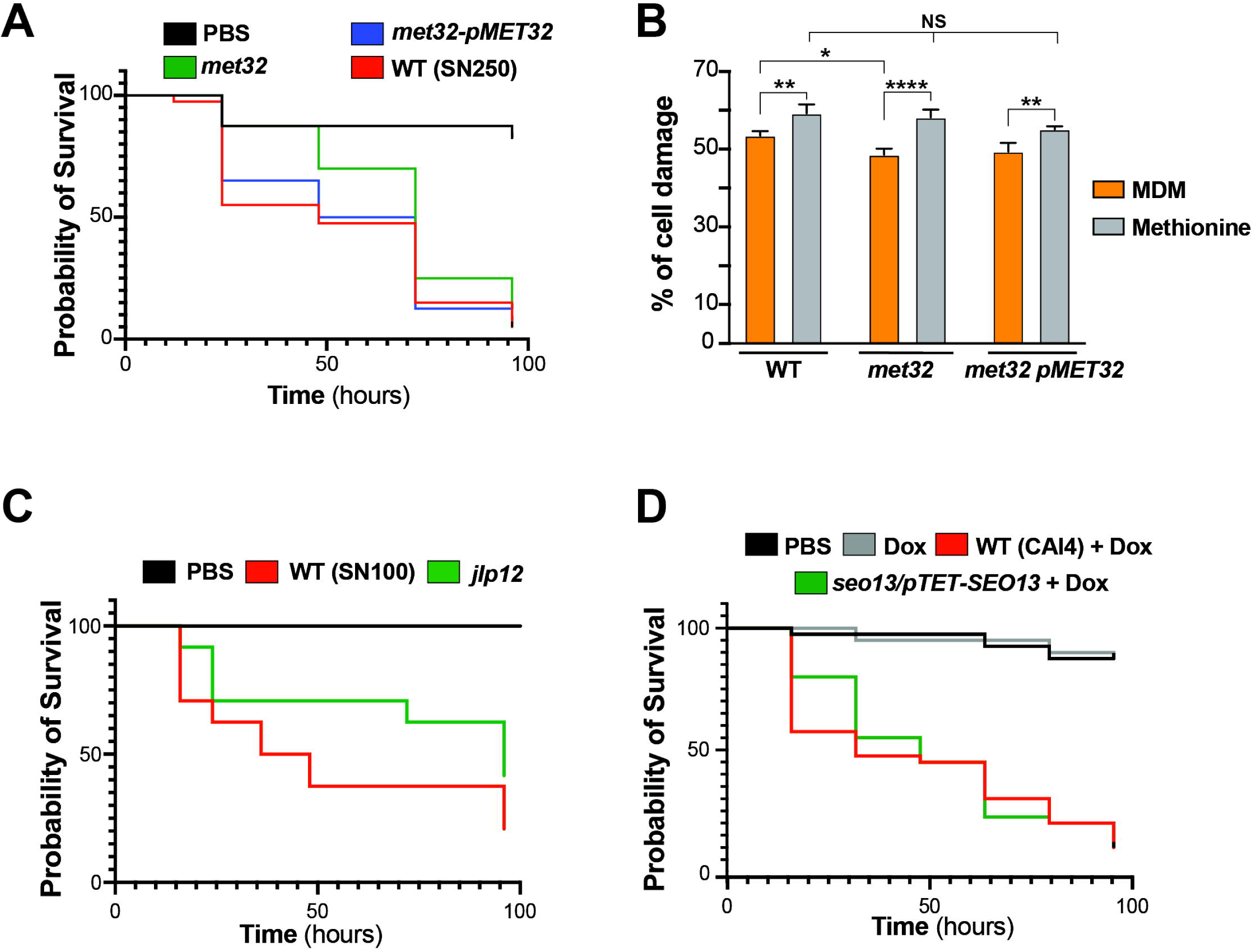
Sulfur utilization processes are required for full virulence of *C. albicans*. (**A-B**) The *met32* mutant exhibits attenuated virulence in *Galleria* model of systemic infection (**A**) and the human enterocyte damage assay (**B**). *Galleria* larvae were injected with WT (SN250), *met32* and the *met32*-p*MET32* complemented strain and survival was monitored at the indicated times for a period of 4 days (**A**). Damage of the human intestinal epithelial cells HT-29 infected with the indicated strains, pre-grown in either MDM or MDM supplemented with methionine (Met), was assessed using the LDH release assay. Cell damage was calculated as the percentage of LDH activity relative to the control condition (HT-29 damaged with 1% Triton X-100) (**B**). (**C-D**) Evaluation of Jlp12 and Seo13 contributions to fungal virulence using the *Galleria* model of systemic infection. Indicated strains were injected to *Galleria* larvae and survival was monitored for 4 days. For the *seo13*/p*TET*-*SEO13* conditional mutant, larvae were first injected with the indicated strains, followed by a second injection of 20 μg/ml of doxycycline (**D**).

We also tested whether inactivation of Met32 effectors, including the sulfonate dioxygenase *JLP12* and the MFS transporter *SEO13*, using the *Galleria* infection model. A similar virulence attenuation was observed for *jlp12* mutant as shown for *met32* (**Figure 7C**). However, for the *seo13* conditional mutant, a slight reduction in virulence was observed only between 15- and 30-hours post-infection (**Figure 7D**). Collectively, these finding suggest that alternate sulfur scavenging promotes virulence of *C. albicans*.

## Discussion

Inside the human host, the opportunistic fungus *C. albicans* thrives in different anatomical sites with contrasting composition and abundance of sulfur-containing metabolites. For instance, in the gut, which is the primary reservoir of *C. albicans*, sulfur pools include hundreds of either organic or inorganic compounds reflecting a vast diversity of sulfur substrates that this yeast could utilize (47, 48). Conversely, diverse microbial residents of the human gut generate elevated levels of RSS, such as hydrogen sulfide, which was shown to exert antifungal activity (45, 49). Thus, to sustain its growth, *C. albicans* and other fungal pathobionts likely employ a versatile metabolic machinery to take benefit from the diversified sources of sulfur sources of the occupied niches. Despite the large studies on *C. albicans* metabolism, sulfur utilization processes and their regulation in this highly prevalent human pathogen remain only partially characterized. The current study comprehensively uncovers biological processes that are modulated by sulfur availability in the opportunistic yeast *C. albicans* and additionally underlines the central role of the transcription factor Met32 in the control of sulfur utilization genes and its contribution to fungal virulence.

The rationale for using different sulfur forms in our RNA-seq comparisons under the repleted-conditions, is to assess whether a specific sulfur utilization pathway is activated to metabolise the corresponding sulfur source. We found that the majority of differentially modulated transcripts were shared among the three sulfur sources (67 % for Met/Cys, 63 % for (NH_4_)_2_SO_4_ and 85 % for taurine; **Figure 1A**) and were enriched in highly similar biological processes (**Figure 1D**). In terms of sulfur metabolism, *C. albicans* upregulated indiscriminately transcripts involved in methionine and cysteine biosynthesis, as well as genes associated with the uptake and utilization of alternative sulfur sources. This reflects that *C. albicans* cells responded to sulfur starvation by activating a broad and non-discriminative response toward a specific sulfur source. Interestingly, this recapitulates the sophisticated carbon metabolic flexibility in *C. albicans*, where simultaneous utilization of diverse carbon sources occurs which contributes to the reinforcement of fungal fitness in nutritionally diversified niches (50, 51). Accordingly, this suggests that *C. albicans* has evolved a robust metabolic machinery to assimilate sulfur-containing metabolites present in mixtures across different niches. The robustness of sulfur utilization in *C. albicans* is also reflected by gene family expansion of many desulfonating oxygenase enzymes as compared to the budding yeast *S. cerevisiae*. For instance, *C. albicans* has five α-ketoglutarate dioxygenases (*JLP*s), seven FMNH_2_-dependent monooxygenases (*FMO*s) and two alkanesulfonate monooxygenases (*SOX*s), while *S. cerevisiae* has one member for each. However, although these enzymes were highly modulated in our RNA-seq dataset, except for Jlp12, inactivation of their respective gene did not result in a growth defect when their mutants were tested for the utilization of various sulfonate substrates. This reflects a functional redundancy between enzymes of the same family and suggests a genetic backup for sulfur utilization processes in *C. albicans*.

In fungi, GSH uptake is mediated by members of the oligopeptide transporters family including Hgt1 in *S. cerevisiae* (52) and Opt7 in *C. albicans* (39). Here, we report first evidence of GSH internalization by a transporter that belongs to the MFS family in fungi. Interestingly, this feature is shared with the pathogenic Gram-negative proteobacterium *Francisella novicida* that exhibits sulfur amino acid auxotrophy and utilizes host GSH as a source of organic sulfur (52). Unlike other bacteria that rely on ABC transporter systems, *F. novicida* imports GSH by the MFS transporter NgtA, which has no significant sequence similarity to the *C. albicans* Seo13. The versatile GSH uptake in *C. albicans* by both the oligopeptide and the MFS transporters might reflect an evolutionary advantage to outcompete microbial cohabitants for GSH. Furthermore, overexpressing *SEO13* partially alleviates the growth defect of *met32* mutant in SLD medium with GSH as the sole sulfur source. This reflects that the control of GSH uptake by Met32 might involve other redundant transporter systems yet to be identified. Of note, Seo13 exhibits sequence similarity (49% homology and 29% identity) with *S. cerevisiae* ScSeo1 that is a member of the allantoate transporter subfamily of MFS shown to be involved in the uptake of methionine analogs (53). While Met32 occupy the promoters of both Seo13 and Seo1, that is the ortholog of ScSeo1, our data suggest that only Seo13 seems to be required for GSH uptake. As *SEO1* was significantly upregulated under sulfur starvation in a Met32-dependent manner, one might speculate that this transporter might mediate the uptake of other sulfur-containing metabolites such as methionine sulphoxide or ethionine as its ortholog in *S. cerevisiae* (53).

Sulfur metabolism leads to the production of different endogenous RSS such as sulfite and sulfide especially during the reductive sulfate assimilation (13). Furthermore, desulfonating enzymes such as α-ketoglutarate dioxygenases (Jlps), degrade sulfonates to sulfite, which will enter the sulfite reduction pathway to generate sulfide (17, 54). At high intracellular concentrations, RSS are toxic and inhibit cytochrome c oxidase and mitochondrial respiration in a manner similar to cyanide (55). Thus, it is not surprising that Met32 coordinate both the control of the utilization of sulfonate and the Sod1-mediated detoxification of RSS catabolites. A similar mechanism was previously described for the *C. albicans* transcription factor Zcf2, which under excess of cysteine, activates parallelly the expression of both the cysteine dioxygenase Cdg1 that catabolizes cysteine to sulfite, and the sulfite efflux pump Ssu1 (44). So far, the unique known RSS detoxification pathway in *C. albicans* is mediated by the Zcf2-Ssu1 module (44, 45). Here, we uncovered a new parallel RSS detoxification pathway which relies on the Cu- and Zn-containing superoxide dismutase Sod1 that is also essential for *C. albicans* to survive oxidative stress and to infect the host (56). The fact that Met32 was required to sustain the basal transcriptional level of Sod1 reflects a housekeeping role of Met32-Sod1 axis in monitoring and neutralizing RSS originated from different endogenous sulfur metabolic routes. In budding yeast and human cells, Sod1 homologs were found to processes a sulfite and a sulfide oxidase activity converting RSS to sulfate (46, 57). Future studies are needed to clarify whether *C. albicans* Sod1 has a similar sulfite oxidase activity that promotes RSS neutralization.

Transcription regulation of methionine and cysteine biosynthesis has been widely studied in both filamentous fungi and yeasts. Overall, this regulation involves a complex network of five transcription factors in *S. cerevisiae* namely Met4, Met32, Met31, Met28, and Cbf1, while filamentous fungi rely only on a single regulator, that is a Met4 homolog (18, 23). Recent work in *C. albicans* underscored that the regulation of methionine biosynthesis is mediated by Cbf1 and Met4, but does not depend on Met28 and Met32, reflecting a case of transcriptional rewiring between the two yeasts (26). A similar scenario was reported in *C. parapsilosis*, with the exception that Met28 function remains associated with methionine biosynthesis as in *S. cerevisiae* (58). Our genome-wide occupancy and RNA-seq data confirmed that Met32 is not associated with *MET* genes while underscoring its role in modulating genes of sulfur uptake and degradation. A similar role has been attributed to Met32 in *C. parapsilosis* based on RNA-seq analysis of *met32* mutant (58). Together, this suggests that in the context of the gradual transition from single protein regulation to a complex circuit of sulfur metabolism, Met32 has specialized in sulfur utilization in the *Candida* CTG clade while it governs methionine biosynthesis in the *S. cerevisiae* lineage. However, the role of the *S. cerevisiae* Met32 in sulfur assimilation could not be excluded as mutants inactivated in both *MET32* and its paralog *MET31* genes, but not *MET32* alone, failed to utilize different alternative sulfur sources (59). Interestingly, we found that both *cbf1* and *met4* mutants showed a sulfur utilization defect as in *met32* (**Figure S4**). This reflects that these regulators might together coordinate the expression of sulfur utilization genes. Cbf1 was found to bind Met32 promoter (27), while our ChEC-seq data did not show any binding of Met32 to the Cbf1 promoter. This let to speculate that Cbf1 might act as a higher hierarchy regulator that controls sulfur assimilation through the Met32-Met4 interplay. Future studies will be necessary to understand the nature of the interplay between the three transcription factors and their hierarchy in the *C. albicans* sulfur utilization network.

We found that inactivation of *MET32* led to both attenuated virulence in *Galleria* larvae and decreased damage to human enterocytes. This attenuated virulence might be linked to the loss of both sulfur metabolic flexibility and the ability to synthesize methionine which consequently will alter the *in vivo* fitness. Beyond the fact that sulfur utilization is important for *C. albicans* infectivity, it is also important to explore the role of Met32 in the context of the commensal lifestyle of this yeast, especially in the gastrointestinal tract. In this niche, *C. albicans* and other *Candida spp*. must compete with millions of other microbial residents for sulfur sources and also must resist the toxicity of increased level of sulfide (29). Of note, the gut of patients with inflammatory bowel disease (IBD) has increased sulfide production due to altered faecal microbial community structure (29, 60, 61). As IBD patients are also characterized by a dysbiotic intestinal microflora with an overgrowth of *Candida spp*., it is tempting to assess the therapeutic value of targeting Met32 to modulate fungal communities and contain the fungal-mediated pro-inflammatory immunity (62–64).

## Materials and methods

### Strains, mutant generation and growth assays

*C. albicans* strains and the primers used for mutant generation are listed in **Table S6**. Growth and maintenance of all strains were carried out at 30°C in fresh yeast-peptone-dextrose (YPD) medium supplemented with uridine (2% Bacto-peptone, 1% yeast extract, 2% w/v dextrose, and 50 µg/ml uridine). Sulfur utilization assay was performed using the Sulfur-Lacking Dextrose (SLD; 1.2 g/l Yeast Nitrogen Base (YNB) without amino acids, ammonium sulfate or magnesium, 4 g/l NH4Cl, 0.84 g/l MgCl ·6H O and 20 g/l glucose) liquid medium lacking any sulfur source (35). Prior to the sulfur utilization assay, *C. albicans* strains were pre-cultured overnight in 5 ml Minimal Dextrose Medium (MDM; 1.7g/l yeast nitrogen base without amino acids and ammonium sulfate, 5g/l (NH_4_)_2_SO_4_, 2% w/v glucose, and 50 mg/l uridine) and washed twice with phosphate-buffered saline (PBS). SLD was used for assaying growth on individual sulfur-containing metabolites at a final concentration of 80 µM. Sulfur utilization assays were carried out under normoxic (21% oxygen) and hypoxic conditions (5% oxygen) using the BioTek™ Cytation™ 5 plate reader with constant agitation at 30°C. For the methionine auxotrophy assay, synthetic dextrose medium (SD) containing 1% glucose, 0.17% YNB without amino acids, 2% agar was used (24). The medium was supplemented with 50 µg/ml uridine, adenine (20 µg/ml), histidine (20 µg/ml), lysine (30 µg/ml), tryptophan (20 µg/ml). Depending on the assay, either methionine (20 µg/ml), or cysteine (20 µg/ml), or both were added. Overnight cultures of *C. albicans* were adjusted to an OD_600_ of 2 and 3 µl of serial dilutions were spotted onto agar plates, followed by incubation at 30°C for 24h. The sulfite tolerance growth assays were performed as follows: overnight cultures were grown in SC medium at a pH of 3.5. Cells were then treated with different amounts of freshly prepared sodium bisulfite (NaHSO_3_) and incubated at 30°C in the Cytation 5 plate reader with continuous shaking and readings taken every 10 mins for 48 hours.

Gene inactivation was performed using PCR replacement cassettes amplified from pFA plasmids as previously described (65). Complementation of the *met32* mutant was achieved by amplifying 1 kb upstream of *MET32* along with the entire ORF. The amplified construct was cloned into pDUP3 plasmid (66) and the resulting pDUP3-*MET32* construct was digested by *SfiI* and integrated into the *NEUT5L* genomic site of the *met32* mutant using the lithium acetate transformation procedure (67). Transformants were selected on YPD plates containing 200 μg/mL nourseothricin and integration was verified by PCR. Genetic rescue experiments were performed by cloning ORFs in the CIp-Act plasmid (68) which was linearized with *StuI* prior to transformation into the *met32* mutant.

### RNA-seq and qPCR analysis

*C. albicans* overnight cultures grown in MDM medium were diluted to an OD_600_ of 0.1 in 250 ml of fresh MDM medium and grown at 30°C under agitation (200 rpm) to logarithmic phase (OD_600_ = 0.6). Cultures were then washed three times with sterile water and transferred to different media including SLD, SLD supplemented with 50µM of different sulfur sources: mixture of Methionine/Cysteine (Met/Cys), ammonium sulfate ((NH_4_)_2_SO_4_), taurine and GSH, and incubated at 30°C for 1 hour. For the sulfite stress experiment, overnight cultures of WT (SN250) and *met32* strains in SC medium were diluted to an OD_600_ of 0.1 in 100 ml of fresh SC medium and grown at 30°C under agitation (200 rpm) to early logarithmic phase (OD_600_ = 0.4). Cultures were then either left untreated or exposed to 600µM NaHSO_3_ for 1 hour. For each condition, two biological replicates were considered for both RNA-seq and qPCR analysis. Total RNA was purified using a glass bead lysis in a Biospec Mini-beadbeater 24 and the RNeasy purification kit (Qiagen) as previously described by Tebbji *et al.* (69).

For qPCR, cDNA was generated from 1 µg of total RNA using the High-Capacity cDNA Reverse Transcription Kit (Applied Biosystems) following the manufacturer’s protocol. The reaction mixture was incubated at 25°C for 10 minutes, followed by 37°C for 120 minutes, and then 85°C for 5 minutes. To remove residual RNA, 2 U/ml of RNase H (NEB) was added, and the samples were incubated at 37°C for 20 minutes. qPCR was carried out using the ABI StepOnePlus system with PowerUp SYBR Green Master Mix (Applied Biosystems) for 40 amplification cycles. The thermal cycling conditions included an initial incubation at 50°C for 2 minutes, followed by 95°C for 2 minutes. This was followed by 40 cycles of 95°C for 15 seconds, 56°C for 30 seconds, and 72°C for 1 minute. Comparative ΔΔCt method and Ct values of studied transcripts were normalized against the Ct values of the *ACT1* gene.

Expression analysis by RNA-seq was performed as previously described (70). Briefly, cDNA libraries were prepared using the NEBNext Ultra II RNA Library Prep Kit and sequencing was performed using an Illumina NovaSeq 6000 sequencing system at the Genome Quebec facility (Centre d’expertise et de services, Montreal). cDNA reads were filtered with fastp version 0.23.2 (71) and mapped to the reference genome sequence with STAR version 2.7.9a (72). The number of reads mapping on each of the genes was calculated by featurecounts version 2.0.1 (73). Differential expression of transcripts was calculated with DESeq2 version 1.40.1 (74). Gene ontology (GO) analysis was performed using GO Term Finder of the Candida Genome Database (75). *cis*-regulatory motif enrichment was assessed in the 500 bp-promoter regions of the top upregulated transcripts in *met32* using MEME analysis software (40). All RNA-seq data are available at the GEO database (https://www.ncbi.nlm.nih.gov/geo/) with the accession number GSE294295.

### ChEC-seq analysis

ChEC-seq (Chromatin endogenous cleavage sequencing) analysis was conducted according to the procedure described by Tebbji *et al.* (76). Met32 was first MNase-tagged using a PCR cassette generated from the pFA-MNase-ARG4 plasmid. Overnight cultures of *C. albicans* MNase-tagged and free MNase strains (control strain) were diluted to a starting OD_600_ of 0.1 in 50 ml MDM medium and grown at 30°C to an OD_600_ of 0.8. Cells were then pelleted and washed twice with PBS buffer and grown for an additional hour in SLD medium. Cells were pelleted at 3,000 × g for 5 min and washed three times with 1 ml buffer A (15 mM Tris [pH 7.5], 80 mM KCl, 0.1 mM EGTA, 0.2 mM spermine, 0.5 mM spermidine, one tablet Roche cOmplete EDTA-free mini protease inhibitors, 1 mM phenylmethylsulfonyl fluoride [PMSF]). Pellets were resuspended in 800μl buffer A containing 0.1% digitonin (Sigma) and permeabilized for 10 min at 30°C with shaking. Calcium chloride was then added to a final concentration of 5 mM, and samples were incubated at 30°C for various time points for MNase digestion. 50 µl of 250 mM EGTA was added to the ChEC digestion mixture to abolish the MNase activity. DNA was purified using the MasterPure Yeast DNA Purification Kit (Epicentre) following the manufacturer’s instructions and resuspended in 50µl of 10 mM Tris-HCl buffer (pH 8.0). RNA was removed by treatment with 10 µg of RNase A at 37°C for 20 minutes. Library preparation, next-generation sequencing (NGS), and peak calling were performed as previously described (76). Motif enrichment was assessed in the top 120 high-scoring peaks using MEME-ChIP software. The ChEC-seq raw data are available in the GEO database with the accession number GSE294295.

### GSH quantification

*C. albicans* cells were grown to exponential phase in 50 ml MDM medium after which the cells were washed twice with PBS. Cells were transferred to SLD or SLD medium supplemented with 80 µM GSH and incubated for 3 hours at 30°C with agitation. Cells were then harvested, washed with PBS and resuspended in 500 µl of pre-chilled methanol and 100 µl of 0.1mm glass beads, and lysed in a Biospec Mini 24 bead-beater at 6800 rpm. The cells were then centrifuged at 10,000 rpm for 10 minutes and the supernatants were collected. GSH quantification was performed using GSH/GSSG Ratio Detection Assay Kit (ab205811, Abcam), following the manufacturer’s instructions.

### Galleria virulence assay

*G. mellonella* larvae in the last instar stage of development, weighing 180 ± 10 mg, from Elevages Lisard (Saint-Bruno-de-Montarville, QC, Canada), were used for this study. Overnight cultures of *C. albicans* strains grown in MDM medium were washed twice and diluted in PBS to obtain 2.5 × 10^5^ cells in 20 μl. PBS alone served as control. The inoculum was injected in a 20 μl volume directly to the last left pro-leg. Larvae were incubated at 37 °C and the number of dead larvae was recorded daily for 5 days. Death was determined based on lack of response to touch and the inability of larvae to right themselves. Kaplan-Meier survival curves were plotted and compared with the log-rank test (GraphPad Prism 10).

### HT-29 damage assay

Human colon epithelial cell line HT-29 (ATCC; HTB-38) was grown in 96-well plates as monolayers in McCoy’s 5A medium supplemented with 10% fetal bovine serum (FBS) at 2 × 10^4^ cells per well and incubated at 37°C with 5% CO_2_ for 24 hours. *C. albicans* cells grown overnight in MDM were pre-cultured in MDM or MDM supplemented with methionine for 6 hours prior to infection. Fungal cells were pelleted by centrifugation, resuspended in cell culture medium and co-incubated with HT-29 cells at MOI (multiplicity of infection) host: yeast of 1:2 for 24 hours at 37°C with 5% CO_2_. A volume of 100 μl of the supernatant was removed from each experimental well and damage to HT29 cells was assessed using a lactate dehydrogenase (LDH) Cytotoxicity Detection Kit PLUS (Roche), which measures the release of the LDH enzyme into the growth medium. LDH activity was determined by measuring the absorbance at 490 nm. Cytotoxicity was calculated as follows: % cytotoxicity = [experimental value− low control (untreated cells)] / [high control (lysis buffer)− low control] × 100.

## Acknowledgments

We thank all members of Dr. Sellam’s lab for their valuable comments. We are grateful to Dr. Joachim Morschhäuser (University of Würzburg), Dr. Malcolm Whiteway (Concordia University) and Dr. Alexander Johnson (UCSF) for sharing strains and plasmids. We also thank Dr Jizhou Li (CHUV, University of Lausanne) for his help with the *Galleria* experiments. This work was supported by funds from the Canadian Institutes of Health Research project grant (grant PJT-180256), the Natural Sciences and Engineering Research Council of Canada (NSERC-DG, RGPIN-2020-06375), the Canada Foundation for Innovation and the Montreal Heart Institute foundation to Dr. Adnane Sellam. Anagha C.T. Menon is supported by a Natural Sciences and Engineering Research Council of Canada CREATE PhD scholarship (EvoFunPath program). Dr. Adnane Sellam is supported by a Fonds de recherche du Québec-Santé Senior Salary Awards.

## Data availability

The original contributions presented in this study are included in the article and its supplemental material. All RNA-seq and ChEC-seq data are available in the GEO database (https://www.ncbi.nlm.nih.gov/geo/) under the accession number GSE294295.

## Supplementary Figures

**Figure S1.** Violin plots showing the distributions of average relative expression levels across the different RNA-seq comparisons using the three sulfur-replete source conditions: mixture of methionine and cysteine (Met/Cys), ammonium sulfate and taurine. The number of differentially modulated transcripts are indicated for each experiments.

**Figure S2.** Phylograms and functional domain organization of *C. albicans* desulfonation proteins including five sulfonate/α-ketoglutarate dioxygenases (**A**), two alkanesulfonate monooxygenases (**B**), the arylsulfatase Ays1 (**C**), and seven FMNH_2_-dependent monooxygenases (**D**).

**Figure S3.** (**A**) Transcript levels of the *C. albicans* GSH transporter *OPT7* are not altered in *met32* mutant. Transcript levels of *OPT7* were measured by qPCR and fold-changes were calculated using the comparative Ct method. For each strain, data were normalized relative to the SLD condition. (**B**) Increasing *OPT7* dosage does not rescue the GSH utilization defect of the *met32* mutant. WT (WT-pAct), *met32* and *met32* overexpressing *OPT7* (*met32*-pAct-*OPT7*) were grown in SLD with 80 µM GSH at 30°C. OD_600_ readings were taken every 10 min for 48 hours. (**C**) Heatmap representation of Met32-modulated transcripts related to different biological processes other than sulfur metabolism. Heatmap showing the modulation of sulfur-related transcripts is indicated in Figure 5D.

**Figure S4.** The transcription factors Cbf1 and Met4 are required for sulfur utilization in *C. albicans*. (**B**) WT (CAI4), *cbf1* and *met4* strains were grown in MDM, YPD, SLD alone or supplemented with 80 µM of different sulfur sources (taurine, methionine, benzene sulfonate and dimethyl sulfoxide (DMSO). Cells were grown at 30°C, and OD_600_ readings were taken every 10 min for 48 hours.

## Supplementary Tables

**Table S1.** Raw RNA-seq data of sulfur starvation in *C. albicans*.

**Table S2.** List of differentially expressed transcripts in *C. albicans* WT under sulfur starvation using a 2-log_2_ FC cutoff and a 1% false discovery rate.

**Table S3.** RNA-seq analysis of *met32* mutant using taurine as a sulfur-replete condition. Modulated transcripts in *met32* mutant were identified using a false-discovery rate of 5% and a ± 1.5-log_2_ FC cutoff.

**Table S4.** Significant Met32 DNA-binding peaks identified by ChEC-seq in *C. albicans* under sulfur starvation.

**Table S5.** RNA-seq analysis of *met32* mutant using GSH as a sulfur-replete condition. Differentially expressed transcripts were identified using a false-discovery rate of 5% and a ±1-log_2_ FC cutoff.

**Table S6**. List of primers and strains used in this study.

