## Supplementary figures and images for "Met32 governs transcriptional control of sulfur metabolic flexibility and resistance to reactive sulfur species in the human fungal pathogen *Candida albicans*"

### Figure S1

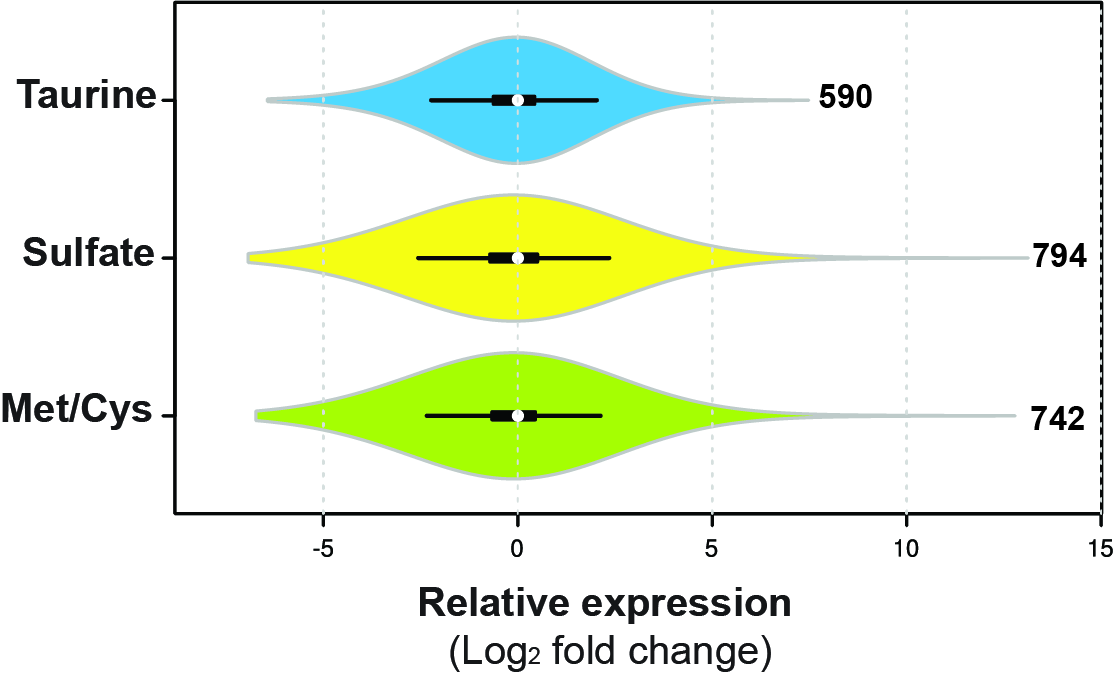

### Figure S2

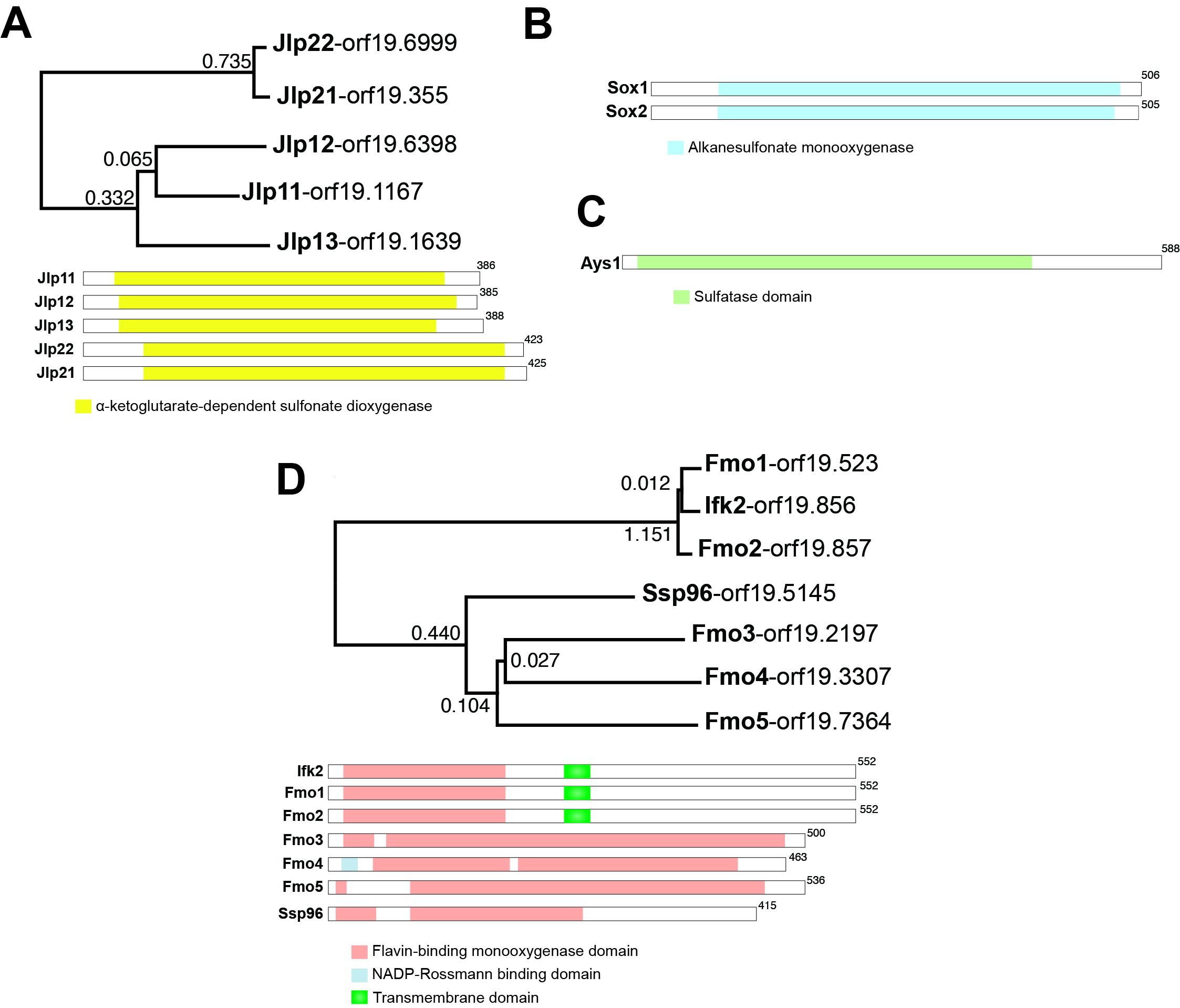

### Figure S3

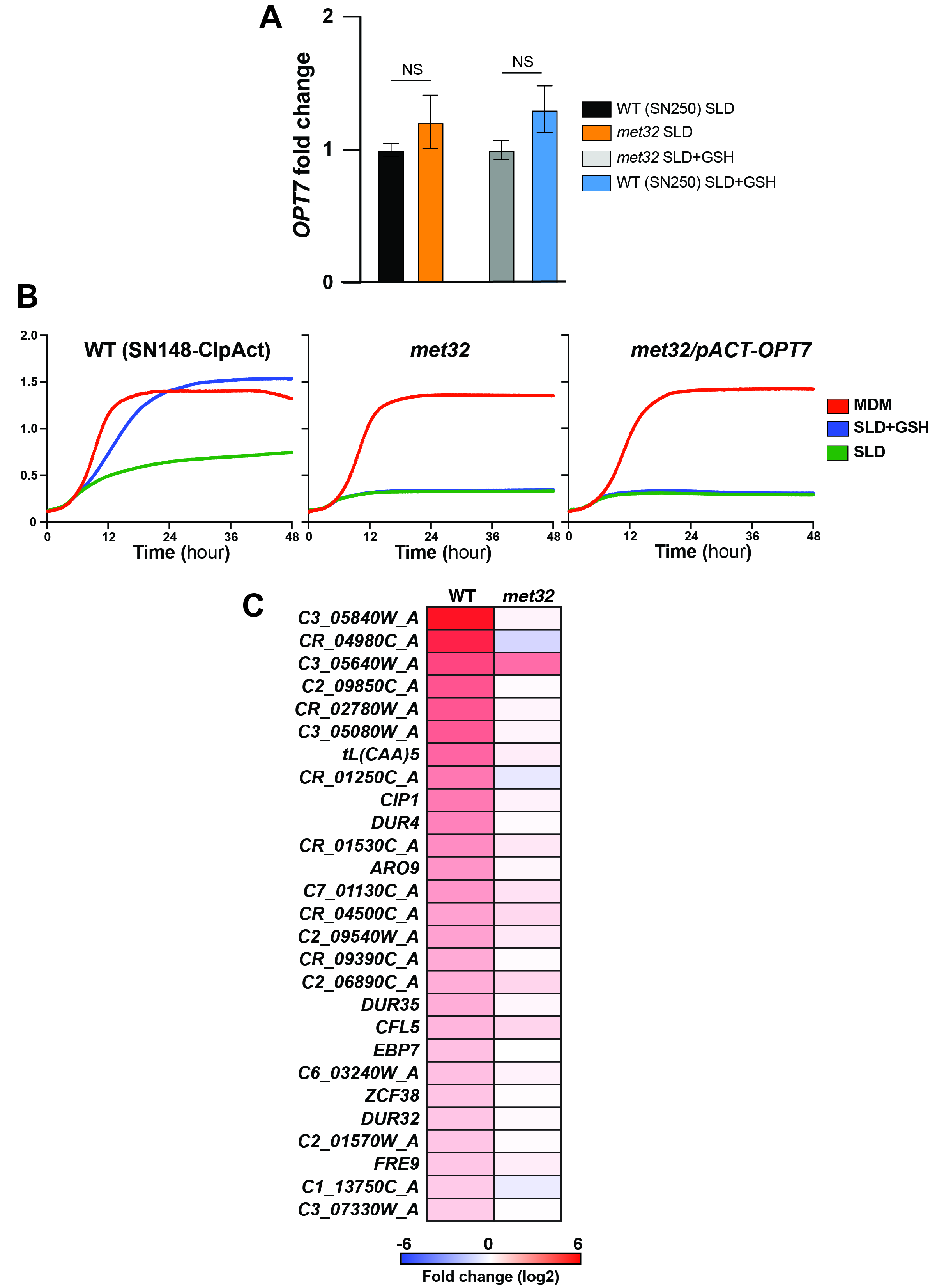

### Figure S4

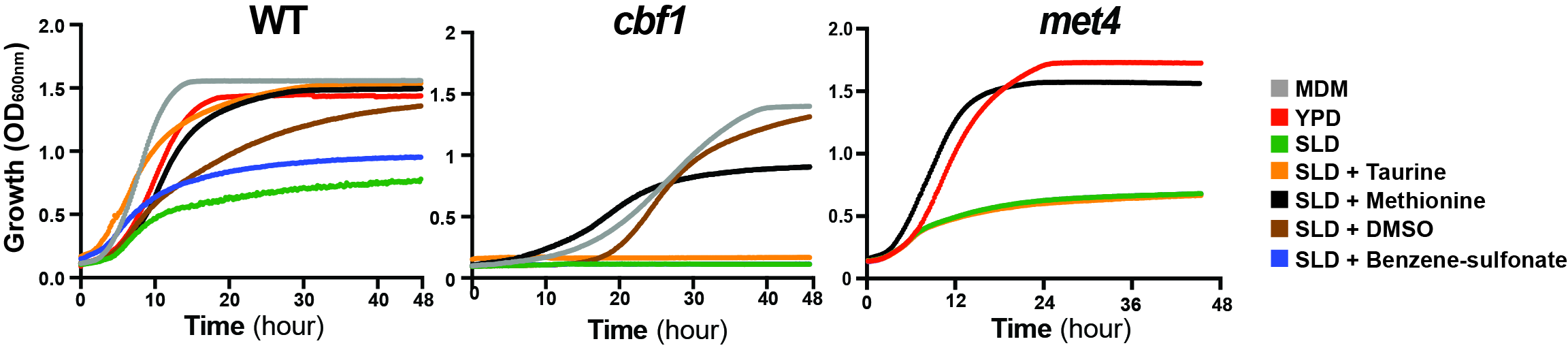
